# Quality-control Normalization of Fluorescence Microscopy Morphometry and Colocalization Measurements for Improved Accuracy and Cross-instrument Reproducibility

**DOI:** 10.1101/2025.10.09.681494

**Authors:** John Oreopoulos, Glyn Nelson, Matthew Gastinger, Christopher L. Morrison, Sabine Thomas, Michael Kiebler, Aidan Boyce, Bernhard Goetze

## Abstract

Quantitative fluorescence microscopy is more reproducible when instrument performance is measured and incorporated into the analysis. We show that routinely monitored quality-control (QC) metrics like the system resolution and inter-channel co-registration are determinant variables that can be used to normalize common image readouts and thereby separate instrument-induced variation from genuine biological changes. For intra-channel morphometry, a Gaussian approximation of the fluorescence imaging process yields analytical factors that predict how geometric measurements (length, separation distance, area, volume, etc.) inflate and scale with resolution blur due to optical misalignments or natural optical quality variations. We validate this behavior by deliberately perturbing the system resolution and by exploiting the natural resolution differences in three nominally equivalent objective lenses configured to image the exact same synapses in cultured hippocampal neurons, where structural differences subtle by eye nonetheless produced statistically significant shifts in measured synaptic puncta volumes. For dual-channel colocalization (overlap) measurements, we normalize the inter-channel co-registration QC metric by the measured point-spread function (PSF) resolutions (rather than theoretical limits associated with the objective lens) and demonstrate how fluorescent pre/postsynaptic cleft protein overlap signals decay in a predictable, exponential fashion as the PSF-normalized registration error increases, with the decay rate depending on the imaged object relative to the PSF size ratio. Mapped field-of-view gradients in channel registration also explain feature orientation flips/rotations and overlap loss without any underlying biological change. Finally, we outline a simple QC-aware microscope normalization workflow where each image measurement dataset is paired with its session PSFs and local co-registration error to remove instrument bias and optionally re-project the results to a declared reference PSF without altering the raw images. This approach improves image measurement accuracy and cross-instrument comparability of experiments and reframes light microscope QC from a passive certification of instrument health into a practical normalization that links the acquisition state to quantitative outcomes, thus ensuring the reliability and interpretability of morphometric and colocalization data in fluorescence microscopy.

## Introduction

Light microscopy is a cornerstone technique of the biological and material sciences, enabling researchers to visualize structures and understand molecular interactions at microscopic scales^1–4^. However, the reliability of the observations and measurements obtained through this technique hinges on quality control (QC) practices^5–12^. Among the many different instrument QC factors that can potentially influence data quality and reproducibility, two critical metrics that come into play in many experimental outcomes are the microscope resolution and chromatic channel co-registration^13–17^. Unchecked variation in these parameters can propagate into image readouts, making comparisons across fields of view (FOV), instruments, and laboratories almost impossible.

Resolution is the ability of the microscope to distinguish between two closely spaced objects and ultimately determines the perceived sharpness of the images^18,19^. Even with the best optical design, this is physically limited by the wavelength of light and the numerical aperture (NA) of the respective objective lens^20^. Typical diffraction-limited performance with a high-NA oil-immersion objective (NA ≈ 1.3–1.4, λ ≈ 520–600 nm) yields lateral resolution on the order of ∼200–250 nm and axial resolution ∼500–800 nm. At these length scales, one can cleanly identify and separate cell bodies, nuclei, mitochondria and lysosomes at the organelle level, trace cytoskeletal tracks and cellular processes such as neurites, and quantify diffraction-limited puncta (e.g., synaptic markers, endocytic pits) as objects; however, sub-organelle features such as synaptic vesicles (40–60 nm), the 20–40 nm synaptic cleft, nuclear-pore substructure, or receptor nanoclusters and similarly sized nanoscopic molecular complexes remain below the resolvable limit and appear blended or enlarged. Paths toward improved resolution include computational deconvolution to restore contrast and moderately sharpen features^21^, structured-illumination microscopy (SIM) achieving ∼100 nm lateral and improved axial resolution^22,23^, stimulated emission depletion (STED) routinely reaching ∼50 nm laterally^24,25^, and single-molecule localization methods (PALM/STORM/DNA-PAINT) achieving ∼10–30 nm precision (with effective resolution set by sampling density and reconstruction)^26–29^. These approaches have revealed previously inaccessible biology such as the layered architecture of focal adhesions^30^, the periodic actin–spectrin lattice in axons^31^, nanoscale receptor nanodomains and their alignment with presynaptic release sites ("nanocolumns")^32^, and the ring-like organization of the nuclear pore complex^33^. Yet each route to superior resolution is accompanied by trade-offs and additional costs, including higher light doses and phototoxicity, demanding labeling density and photophysics, longer acquisitions and drift sensitivity, complex reconstruction/calibration routines, and for some methods, limited compatibility with thick or dynamic live samples.

Chromatic co-registration refers to the 3D overlay accuracy between a microscope’s distinct spectral channels and determines how faithfully spatial relationships and interactions between fluorescently labeled biomolecules or targeted molecular species are represented^34–36^. Departures from ideal overlay arise from both longitudinal (axial focal shifts) and lateral (color-dependent magnification/translational) chromatic aberrations, as well as dispersion mismatch, cover-glass thickness errors, filter tilt/wedge angles, and misalignments in the excitation or detection paths. These image registration shift mechanisms motivate two classes of remedies: hardware designs and alignments that minimize color error at the source, and software corrections that estimate and remove the remaining offsets. Hardware strategies to improve registration include apochromatic or super-apochromatic objectives with extended color correction, matching tube lenses and relay optics to objective design, minimizing wedge/tilt in filters, selecting multi-band dichroics engineered for low lateral color, and using immersion media whose dispersion matches the specimen^37^. Recently, reinforcement learning combined with tunable lenses have also been employed for real-time correction of chromatic aberrations^38^. Careful alignment of excitation and detection paths plus routine microsphere-based checks help maintain both on-axis and off-axis overlay^12^. Software approaches register channels post-acquisition or in real-time using mapped microsphere fields, embedded fiducial markers, and biological landmarks to estimate global rigid transforms (translation/rotation/scale) or spatially varying warps (polynomials, thin-plate splines, piecewise-affine), extended to 3D to handle axial chromatic focal shifts^39–44^. Image cross-correlation correction methods that do not require fiducial markers or mapped microsphere offsets have also been used^45^. When calibrated per instrument and objective, these software approaches can bring channels into sub-pixel agreement across much of the FOV and markedly stabilize two-color measurements. Even so, chromatic corrections have limitations as well: Chromatic shifts often vary with imaging depth and refractive-index mismatch, so a single 2D mapping may fail in thick samples; non-rigid warps can subtly distort morphology, and pixel intensity resampling introduces blur and pixel correlations. Furthermore, transformations of image datasets may not perfectly transfer to heterogeneous specimens, and for live-cell imaging, the added processing can reduce temporal resolution or fail under drifting illumination, lateral stage, and axial focus conditions.

Despite the availability of super-resolution and sophisticated chromatic-correction solutions, conventional, diffraction-limited fluorescence imaging remains the workhorse of cell biology. It is fast, comparatively gentle on live samples, broadly compatible with common labels and thick specimens, inexpensive to acquire and maintain, and straightforward to train across laboratories. It also scales to high-throughput screens and longitudinal studies while preserving continuity with the vast legacy of archival datasets. As light sources, advanced optics, and cameras become cheaper and more uniform across core facilities and labs, the limiting factor for experimental comparability is increasingly practice, not hardware, thus making standardized QC both timely and impactful. Indeed, the continued expansion of the consortium for Quality Assessment and Reproducibility for Instruments and Images in Light Microscopy (QUAREP-LiMi), comprising over 500 members across academia and industry globally, is a sign of the growing recognition of the critical importance of and desire for QC in microscopy related research^46,47^. Without shared and standardized practices, though, variability across instruments and associated protocols will obscure biological effects and limit cross-study comparability. Accordingly, our goal in this paper is to provide an unambiguous, experimentally grounded and theoretically supported demonstration of why routine light-microscope QC is essential. We image the same biological structures while deliberately varying two core instrument metrics— resolution and chromatic co-registration—and show that these QC values predict the direction and magnitude of changes in standard morphometry and colocalization readouts. To make this link explicit and usable, we develop a compact, analytic normalization treatment that connects QC to measurement bias and supply a practical rationale for normalizing conventional fluorescence imaging measurements when those measurements are close in scale to the related QC metrics. To operationalize this motivation, we adopt QUAREP-LiMi–aligned procedures to characterize instrument performance and investigate corresponding consequences for standard readouts^48,49^. Specifically, we quantify resolution along with inter-channel offsets and then re-image matched cellular structures under distinct microscope performance conditions to test the predicted effects on morphometry and colocalization.

As a biologically-grounded model system for linking these two QC metrics to image readouts, we focus on presynaptic and postsynaptic protein puncta in cultured hippocampal neurons. These synaptic compartments have a well-established nanometer-scale organization from electron microscopy^50,51^ and super-resolution light microscopy studies^52–54^, yet, when imaged with conventional, diffraction-limited light microscopy^55^, they exist at the edge of optical resolution. In excitatory synapses, the active zone (AZ) and postsynaptic density (PSD) face one another across a ∼20–40 nm cleft. Commonly used antibody epitopes (e.g., vGLUT, Syanpasin, and Bassoon at the AZ and Homer/PSD-95 at the PSD) are offset in 3D by roughly 120–160 nm. Consequently, two-color fluorescence microscopy image readouts that appear coincident at the pixel level often reflect tightly apposed but non-identical structures—exactly within the geometric regime where traditional light microscopy resolution and chromatic co-registration most strongly modulate measured volumes, overlaps, and pixel-wise coefficients. Image assays of these neuronal structures are also methodologically mature and widely adopted. For example, fixed-cell synaptogenesis workflows typically immunolabel a presynaptic marker (e.g., Bassoon) and a postsynaptic marker (e.g., Homer or PSD-95) and then quantify either (i) punctum volume/intensity per channel or (ii) pre/post colocalization (overlap fractions, object-pair counts, or pixel-wise coefficients). Popular pipelines such as the ImageJ Puncta Analyzer define a synapse as an apposed pre/post pair (often with an “overlap-pixel” or sub-diffraction centroid-distance rule), making the readout both standardized and conveniently re- analyzable under different QC regimes^56^. Biologically, these structural metrics are meaningful. Puncta volume and intensity correlate well with synaptic organization and function (e.g., AZ scaffold size and release probability presynaptically; PSD scaffold/receptor content postsynaptically)^57^, while colocalization-based synapse counts track developmental, activity-dependent, and disease-related changes^58^. Many groups pair the structural readouts with functional measurements (mEPSC frequency/amplitude, pHluorin/iGluSnFR/GCaMP reporters) or ultrastructural validation, reinforcing their interpretability^59,60^. Critically for this study, the same features that make synapses biologically interesting also make them QC-sensitive. Because true pre/post separations are sub-diffraction, yet non-zero in size, even modest changes in the PSF (resolution) can inflate apparent punctum volumes and artificially increase overlap, whereas tens-of-nanometers of chromatic mis-registration can erase true appositions or create spurious false-positive ones. Leveraging instrument-derived QC maps, we show that these biases predictably reshape common observations of synaptic morphology across the FOV and across objective lenses, enabling principled normalization and improved experimental design. Although we illustrate the analytic formalism and companion workflow for such observations, the same QC-to-measurement relationship applies to any image assay where feature sizes and separations are on the order of the system’s resolution.

## Methods

### Notation

Throughout this work, uppercase axis labels (X, Y, Z) are used to denote the physical axes of the microscope hardware (e.g., motorized XY stage translations or axial Z-scans), whereas lowercase symbols (x, y, z) are used for spatial coordinates in the image and analysis domain. This convention distinguishes instrument motions from mathematical variables. In the present study, the instrument and coordinate systems are aligned such that X, Y, and Z correspond directly to x, y, and z, respectively. In particular, the Z-axis of the focus drive coincides with the optical axis of the imaging system.

### Microscope equipment

All experiments were carried out on an Andor Technology / Oxford Instruments Benchtop Confocal Microscope (BC43) equipped with three different plan-apochromat 60× oil immersion objective lenses having nearly the same NA, but were of different optical design and model generation (Objective 1: Nikon 1.42 NA CFI Plan Apochromat Lambda D series, Model# MRD71670; Objective 2: Nikon Plan Apochromat XA 1.40 NA, model# MRD01600; Objective 3: Nikon 1.40 NA CFI Plan Apochromat Lambda series, Model# MRD01605). This spinning-disk confocal microscope^61,62^ uses a 4.2 megapixel (2048×2048 pixels) scientific complementary metal-oxide-semiconductor (sCMOS) camera detector having 6.5 μm×6.5 μm size pixel elements. At 60× magnification, the back-projected sample plane pixel size was almost exactly 100×100 nm and was confirmed by imaging the “field of rings pattern” of the Argolight Argo-HM calibration slide^63^ (Argolight SA, France). The BC43 has four distinct spectral channels utilizing a common multi-band dichroic mirror in combination with directly modulated lasers for fluorescence excitation and bandpass interference filters loaded in a rotating filter wheel for viewing fluorescence emission. Only the “Green” (488 nm excitation; 521/24 nm emission) and “Yellow” (561 nm excitation; 593/31 nm emission) channels were used for this study. All images were acquired using the confocal imaging mode of the microscope (50 μm pinhole physical size on the disk). The same immersion oil was used for all three objective lenses at room temperature (Thorlabs OILCL30; Cargille Type LDF with refractive index 1.518).

### Neuronal cell culture

Cell cultures from primary rat hippocampal neurons were generated as previously described^64^. In short, embryos of timed pregnant Sprague-Dawley rats (Charles River Laboratories) were used to isolate hippocampi of embryonic day 17 (E17). Cells were dissociated and plated on Poly-L-Lysine coated cover slips and cultured in NMEM+B27 supplemented medium (Invitrogen). Immunostaining was performed with cultured neurons on day 17 in vitro (DIV).

### Immunostaining and sample preparation

Neurons were fixed for 15 minutes with 4% paraformaldehyde (PFA) and immunostained as previously described^65^. The following primary antibodies were used for incubation at 4°C overnight: polyclonal guinea pig anti-vGLUT 1:500 dilution (Synaptic Systems) and polyclonal mouse anti-Homer 1:500 dilution (Synaptic Systems). On the next day, the following secondary antibodies were incubated for 1 hour at room temperature: anti-guinea pig AlexaFluor488 and anti-mouse Cy3 conjugated antibodies 1:1,000 dilution (Life Technologies). DAPI staining was carried out for 5 minutes at room temperature. Coverslips were mounted on microscope slides with ProLong Diamond Mounting Medium (Life Technologies).

### Quality control and instrument characterization

The most common and practical method for experimentally measuring the microscope PSF involves imaging sub-resolution fluorescent microspheres^13,12,48^. The microspheres act as idealized point sources of fluorescence emission and their image, when captured by the microscope, directly represents the system’s PSF. In a well-aligned system using high-quality objective lenses and optics, the PSF response near the middle of the imaged FOV is constant (isoplanatic), thereby producing the most reliable imaging results. However, even in systems such as these, it is not uncommon to observe deviations from the central, on-axis PSF response in the image corners as well as at greater depths within inhomogeneous samples^66^. The microscope’s channel pair-dependent chromatic co-registration can also be measured in a similar manner using multi-color emitting microspheres and localizing the 3D center of the microsphere in each spectral channel to measure the positional offset of different wavelengths^49^. Similarly to how the microscope’s PSF can exhibit a spatial dependence across the FOV, so too will there often be differences in channel registration across the FOV, and these spatial variations in channel overlap can considerably impact upon measures of co-localization or overlap between two fluorescently labelled biomolecules.

Since this report attempts to directly and unambiguously demonstrate how changes in a light microscope’s system QC metrics can have a direct impact on experimental outcomes, stringent efforts were made to image the exact same FOV within all examined samples with each objective lens using identical laser powers and exposure times while simultaneously limiting and minimizing light exposure to avoid specimen photobleaching. Before imaging, the microscope was allowed to reach thermal equilibrium, ensuring negligible axial (Z) drift during data acquisition. Sequential dual-channel 3D confocal Z-stacks (10 μm Z-range, 100 nm Z-step) of 200 nm multi-color TetraSpeck fluorescent microspheres mounted in Saint Gobain BC-600 Optical Cement clear epoxy resin (refractive index 1.56) between a 170 μm thick glass coverslip - 1 mm glass slide sandwich (Thermo-Fisher T14792) were acquired. Acquisition FOVs were carefully selected for those exhibiting well-dispersed single microspheres on the coverslip surface with good coverage right to the image FOV corners and avoiding FOVs possessing obviously bright microspheres (which were likely to be aggregates). Once selected, the same FOV was scanned with each 60× objective lens to ensure that the characterization of each objective in use with the BC43 microscope would be completely fair and equal. Multiple FOVs were imaged in the same manner with each objective lens in sequence as replicates to confirm the repeatability of the image measurements taken with the first FOV. These image datasets were subsequently used to create quantified spatial heatmaps of the (Green channel) resolution and (Green-to-Yellow channel) chromatic co-registration of each objective lens (see below for more details). Z-stacks of the Argolight Argo-HM slide “field of rings” pattern were also acquired with each objective lens using the same acquisition settings as an alternative sample type to assess and confirm measurements of chromatic co-registration made using the TetraSpeck microsphere sample.

Full FOV lateral (XY) and axial (Z) heatmaps of the system PSF in conjunction with each 60× objective lens were created using the freely available and open-source software package PSFj^14^. Very briefly, this program processes 3D Z-stacks of fluorescent microspheres by first automatically identifying individual microspheres through size and intensity thresholds and then subsequently fitting a 3D Gaussian ellipsoid to each microsphere using non-linear regression techniques. After accounting for the sample and immersion medium refractive index, as well as the voxel size and microsphere size (100 nm x 100 nm x 100 nm and 200 nm diameter respectively in our case), PSFj extracts the minimum and maximum lateral full-width at half- maximum (FWHM_min_ and FWHM_max_ respectively) as well as the axial full-width at half-maximum (FWHM_z_) Gaussian model fit parameters associated with each identified microsphere. Since the Airy disk diffraction pattern of a fluorescent point-source object is well approximated near its center by a Gaussian function, the fitted Gaussian widths (FWHM) are commonly used as estimates of the lateral and axial resolution^12,19,48^ (see also Supplementary Text). For this study, we ignored the FWHM_min_ values and instead focused on the more conservative FWHM_max_ values to denote the measured lateral resolution. Having also measured and tabulated the 3D sub-voxel localization coordinates of each identified microsphere, the program then outputs color-coded, continuous heatmaps of the lateral and axial resolution using a sliding-window median filter that assigns each heatmap pixel the local median FWHM value of nearby microspheres. PSFj also possesses a dual-channel analysis mode which can conduct the same image analysis pipeline on image datasets of multi-color microspheres and can create dual-channel colored heatmaps of the lateral and axial chromatic mis-registration by computing the lateral and axial separation distances between identical microspheres seen in both channel datasets. For single-channel and dual-channel analysis modes, PSFj also outputs a full spreadsheet file listing the found coordinates of each processed microsphere in both channels along with the measured lateral and axial FWHM values. A custom-written Python programming language-based executable application (see download link in Supplementary Text) was used to convert the listed dual-channel microsphere sub-voxel localized coordinates into measured lateral and axial chromatic separation distances and then subsequently transformed into linearly interpolated co-registration heatmaps according to the definition of co-registration set out by Faklaris *et al.*^12^. Just like with the resolution heatmaps, interpolated spatial representations of a microscope system’s co-registration like this allow for easy look-up of where within the FOV the inter-channel 3D mis-registration is minimal and to what quantifiable degree. We also normalized the spreadsheet FWHM_max/z_ values against the theoretical lateral/axial resolutions for each objective lens and replotted them against each other on 2D scatter graphs, as was demonstrated by Faklaris *et al.*^12^. Lateral and axial channel separation distances were normalized by the *measured* lateral and axial resolutions and similarly plotted against each other. Transformations of the PSFj data output like this permit a quick and easy comparison of system resolution and co-registration across different microscope instruments (or equivalently in our case, the same system using different objective lenses).

### Cellular imaging

The cultured neurons were imaged in the same manner as the QC calibration samples. Sequential dual-channel (Green; Alexa488-vGLUT and Yellow; Cy3-Homer) 3D confocal Z-scans of the fluorescent pre- and postsynaptic proteins in the neuronal cells were captured with all three objective lenses consecutively, again taking great care to maintain the laser power and exposure time acquisition settings within the exact same sample FOV each time. For some cells, after acquiring the initial Z-scan, the sample was immediately translated with the motorized XY-stage such that a dendrite of interest in the middle of the FOV was moved to the upper left-hand corner of the FOV and another Z-scan was acquired. For these cells imaged in the middle and corner of the FOV, the same procedure was also immediately followed with the other two objective lenses. These paired middle and corner FOV-dependent datasets were used for the co-registration studies described in the results section.

### Neuronal cell image analysis

All 3D surface objects were created using Imaris 10.2 (Oxford Instruments). The creation parameters were constructed using the built-in Imaris machine learning pixel segmentation method. To provide a consistent and reproducible surface creation, the detection algorithm was trained on the same puncta from all 3 objective lenses simultaneously. Training pixels within the XY and XZ planes yielded accurate detection of the fluorescent edges of the puncta. This training process was followed independently on both the Green and Yellow channels. Each successfully identified Yellow channel puncta was paired with a corresponding Green channel puncta. Following object creation, the resulting surfaces were filtered by intensity, shape, and size. For each objective lens and region of interest (ROI), the same puncta were rendered and analyzed. Inter-channel, (object-based) colocalized puncta volume overlap was defined by measuring the percentage overlap enclosed between the pre- and postsynaptic puncta surfaces while colocalization was calculated using the traditional Pearson’s correlation coefficient (PCC) definition^15^.

### Operational definition of puncta (vGLUT and Homer)

In this study, a *punctum* denotes a contiguous 3D fluorescence object segmented from a single channel’s voxels after background correction, thresholding, and size filtering. When the Green channel reports a successful segmentation (vGLUT), the resulting punctum represents a single presynaptic site at light microscope resolution—i.e., the AZ scaffold and/or its associated vesicle-rich cluster, not an individual vesicle or molecule. In contrast, when the Yelllow channel reports a segmentation (Homer), the resulting punctum is the local postsynaptic specialization— i.e., the excitatory (PSD) or inhibitory (scaffold/receptor) density, not an individual receptor complex. In all image analyses, “puncta” refer to these segmented 3D masks. Volume and overlap/colocalization image measurements were computed directly from the associated binary masks and intensity counts within their voxel supports.

## Results

### How resolution impacts sub-cellular volume image measurements

We designed two experiments based around one microscope stand to directly demonstrate the effect of relatively minor optical misalignments and component alterations on intra-channel morphometric image measurements. In the first experiment, acting as a positive control, we purposefully reduced the effective NA of Objective 1 by constricting the back-aperture of the objective lens with an internal iris positioned near an equivalent/conjugate back-focal plane in the fluorescence emission pathway relay optics (Fig. 1). This motorized iris normally opens and closes automatically via firmware settings when the image mode is toggled between fluorescence and differential phase contrast^67^ (DPC) modes to suppress background noise and enhance contrast respectively. The aim of these artificially induced resolution changes was to be deliberately subtle with the system perturbations to mimic real world scenarios where a microscope may well have some minor aberration or misalignment error in the light path, but the user would not necessarily notice through visual inspection of the images unless they had performed a quantitative QC check. This restriction of the objective back aperture in the emission pathway was expected to increase the lateral and axial FWHM compared to the Full NA unrestricted condition, and indeed we measured a 12% lateral and 18% axial widening of the PSF associated with a single microsphere located in the middle of the FOV (ie: reduced resolution) in the most restricted condition (Restricted NA 2, see Fig. 2).

**Fig 1.**
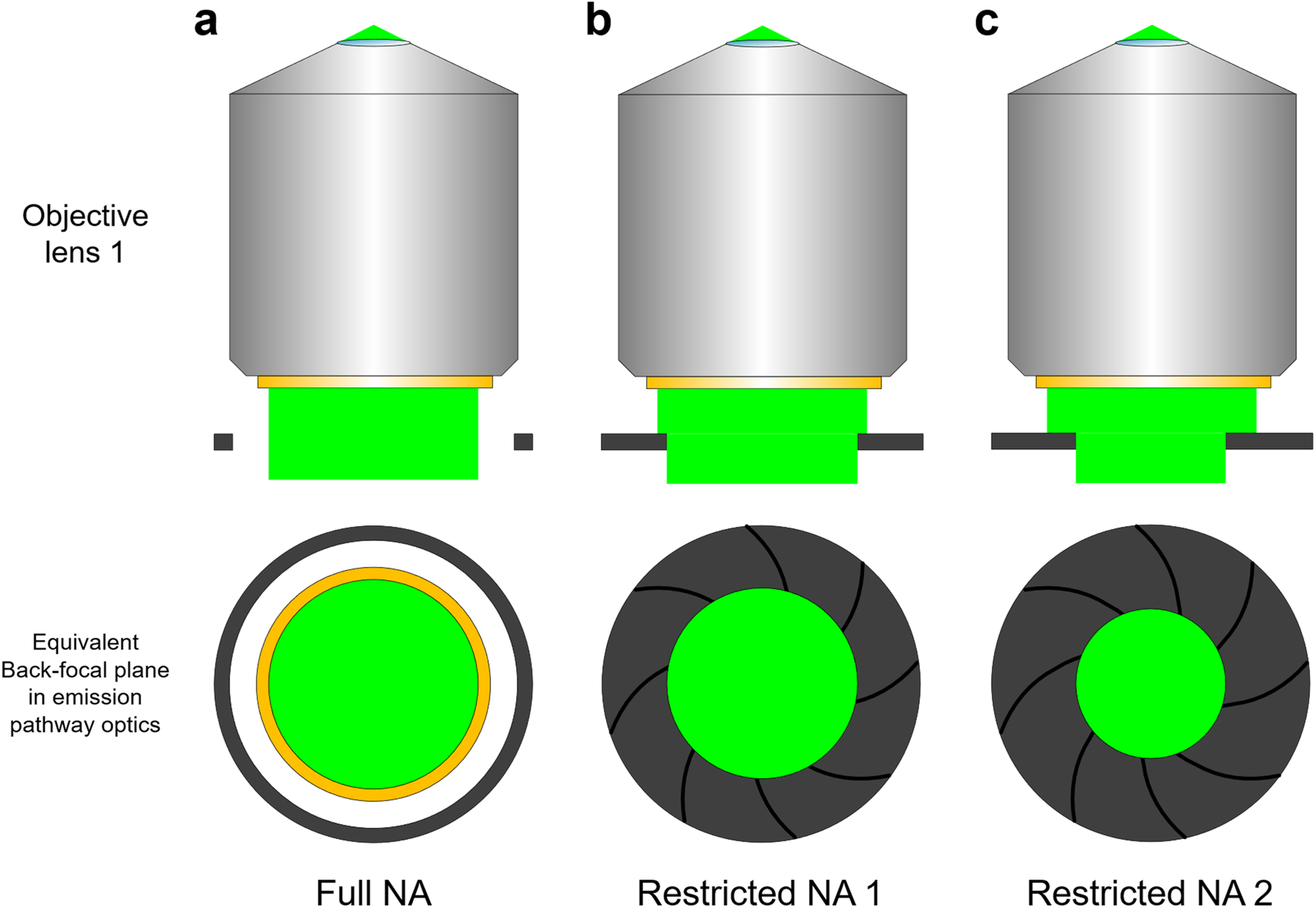
Schematic representation of the resolution change positive control experiment. Fluorescence images of QC microsphere calibration samples and fluorescent neurons were captured with Objective 1 using an increasingly restricted emission pathway beam through means of a motorized iris positioned in an equivalent back-focal plane of the microscope optical train. Radial constriction of the emission beam diameter in this infinity space is equivalent to reducing the effective emission numerical aperture associated with the objective lens and therefore should lead to a reduction in image resolution as judged by the PSF extent of the microspheres or the volumes of small dendritic spine puncta in the neurons. **a,** The Full NA of the objective lens is utilized to image the sample since the completely open iris exceeds the beam size and back aperture diameter of the objective. **b,** A slightly closed iris position (Restricted NA 1) vignettes the periphery of the emission beam and induces a small, but measurable resolution degradation. **c,** The iris was closed down further (Restricted NA 2) to make the resolution degradation more obvious and greater in magnitude. Schematic not to scale.

**Fig 2.**
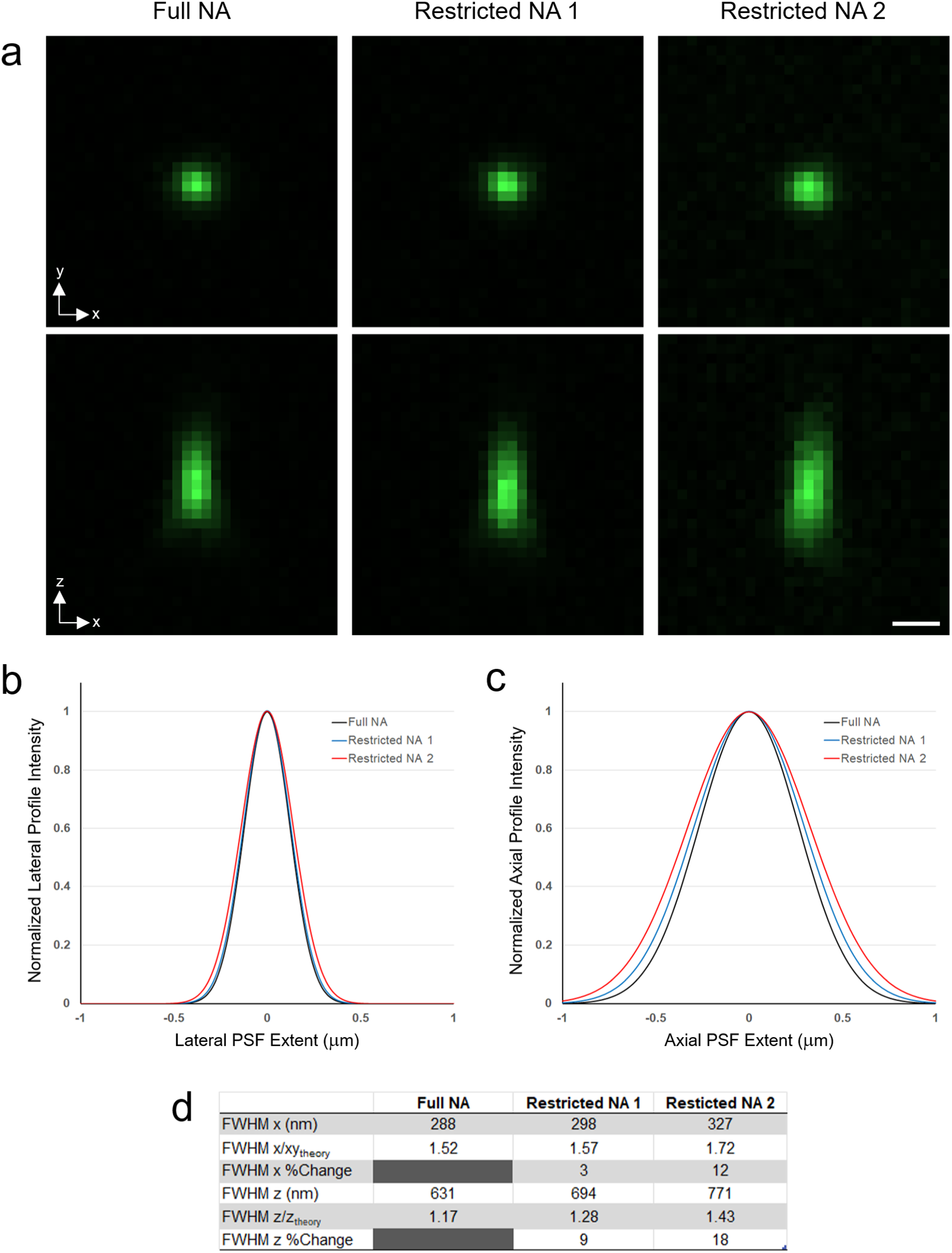
PSF analysis of a single fluorescent microsphere observed with varying emission pathway NA conditions as described in. Fig. 1**. a**, Lateral XY (top row) and axial XZ (bottom row) maximum intensity projections of the microsphere for each emission NA setting. Scale bar = 0.5 μm. **b**, Normalized lateral and, **c**, axial fitted Gaussian intensity line profiles of the microsphere for each NA setting. **d**, The associated absolute and normalized FWHM profile values are tabulated with the relative difference to the Full NA condition.

By eye, the resolution change was subtle and barely measurable under the Restricted NA 1 setting but became more pronounced under Restricted NA 2, especially in the axial (XZ) projection of the microsphere. This behavior was expected since axial resolution scales approximately as 1/NA^2^, whereas lateral resolution scales as 1/NA^19^. Even with the larger degradation at Restricted NA 2, the magnitude of the resolution change remained difficult to estimate by visual inspection alone, underscoring that qualitative impressions can diverge from quantitative PSF-based measurements. This is the first example we show where visual impressions of an image do not correlate well with quantified image measurements.

We then measured the microscope PSF over the entire FOV to obtain a better statistical sampling and look for signs of any spatial dependencies of the system’s resolution (Fig. 3). The average lateral and axial FWHMs for the two restricted NA conditions across the FOV showed significant increase in lateral and axial spread of the PSF (16% and 19% respectively for the most restricted NA (Fig. 3e), again in line with the expected resolution reduction due to a purely reduced detection NA. Generally, the microsphere’s PSF lateral extent uniformly inflated across the FOV because of the induced aberration, whereas the axial extent showed a slight spatial dependency with the PSF broadening, biased mostly in the middle region of the FOV (Fig. 3b and e).

**Fig 3.**
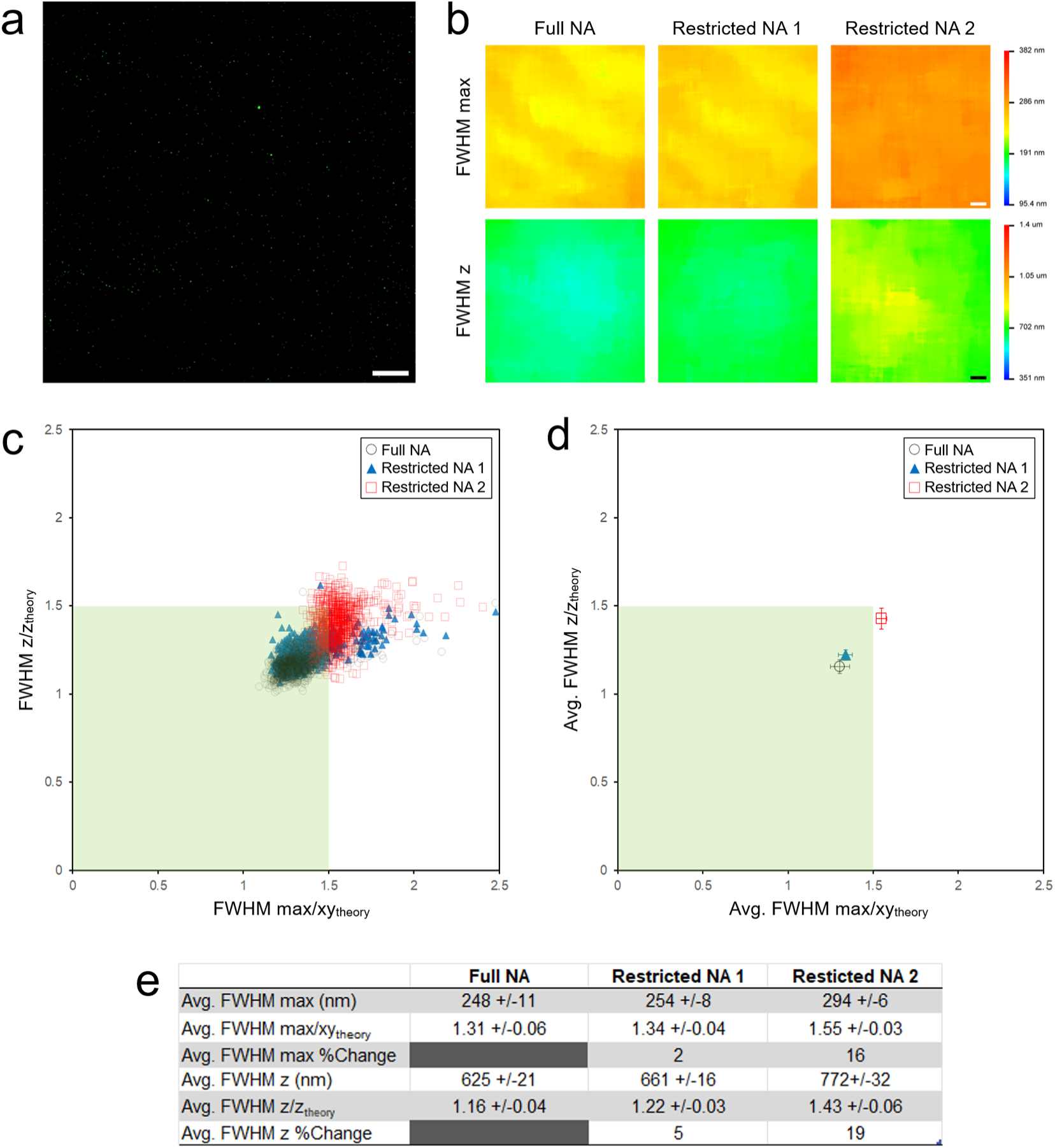
Full FOV microscope lateral and axial resolution characterization of Objective 1 under the varying emission pathway NA conditions. **a**, Green channel maximum intensity Z-projection, full FOV image of 200 nm fluorescent microspheres acquired with Objective 1 at Full NA. **b**, PSFj lateral (FWHM_max_) and axial (FWHM_z_) resolution heatmaps of Objective 1 for each emission pathway iris condition of the resolution control experiment. Scale bar in **a** and **b** is 20 μm. **c**, For each NA setting, the lateral and axial FWHM values for each isolated microsphere in **a** were normalized by their theoretical values for Objective 1 (FWHM_xy,theory_ = 190 nm; FWHM_z,theory_ = 541 nm) and plotted against each other. The green box delimiting the 1.5x theoretical resolution ratios match the recommended microscope resolution performance tolerance limits of Faklaris et al.^12^. **d**, The centroids or each data “cloud” in **c** plotted the same way representing the average resolution performance across the FOV for each NA setting of the control experiment. **e**, Average FOV resolution FWHM measurements in tabular form, with the relative change in relation to the Full NA scenario.

Normalizing the measured FWHM_max_ and FWHM_z_ for each microsphere in the FOV by Objective 1’s Full NA theoretical lateral and axial resolution (FWHM_xy_,_theory_ = 0.51λ_EM_/NA, λ_EM_ = emission wavelength; 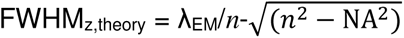, *n* = immersion media refractive index) and plotting these data points against each other on a Cartesian graph produced a “cloud” of lateral/axial resolution pair data points which allowed us to assess the overall resolution behaviors for each of the three NA conditions of the positive control experiment (Fig. 3c). Furthermore, we determined the average resolution behaviors for each of these NA conditions by plotting the centroid of each data cloud (Fig. 3d). With these data transformations, we observed that the resolution changes due to the purposefully “misaligned” emission pathway iris closure was sufficient to push the judged resolution performance of Objective 1 from a level of “acceptable” at the Full NA condition (both FWHM ratios <1.5) to an “unacceptable” level according to the tolerance recommendations of Faklaris *et al.*^12^ and the QUAREP-LiMi standards community at the Restricted NA 2 condition (lateral FWHM_max_/_theory_ ratio >1.5).

An alternative investigation of the PSF’s lateral broadening was performed using Fourier domain (spatial frequency) analysis techniques yielding a similar degraded resolution trend under progressively closed down detection NAs by examining the radially averaged modulation transfer function (MTF) resulting from the Fourier transform of maximum intensity Z-projection images of the microspheres^68–71^ (see Supplementary Fig. 1). Although this complementary method offers a global assessment of resolution changes and reinforces the findings described for lateral resolution, it lacks the 3D spatial context that traditional spatial domain PSF fitting techniques provide.

Having successfully established the changing resolution behaviors observed using the QC microsphere sample, we then carefully imaged dendritic spine presynaptic puncta via fluorescently tagged AZ vGLUT proteins in cultured hippocampal neurons within the exact same FOV for each corresponding NA condition (Fig. 4). As with the QC sample, for this positive control experiment we expected to observe changes to the cell images that would correlate with the degraded resolution of the restricted NA conditions. In particular, since the immunostained neurons contain punctate spatial features which were small and comparable to the resolution limit associated with Objective Lens 1, the volumes of these features should correspondingly enlarge due to the resolution blurring induced by the emission pathway iris closing down.

**Fig 4.**
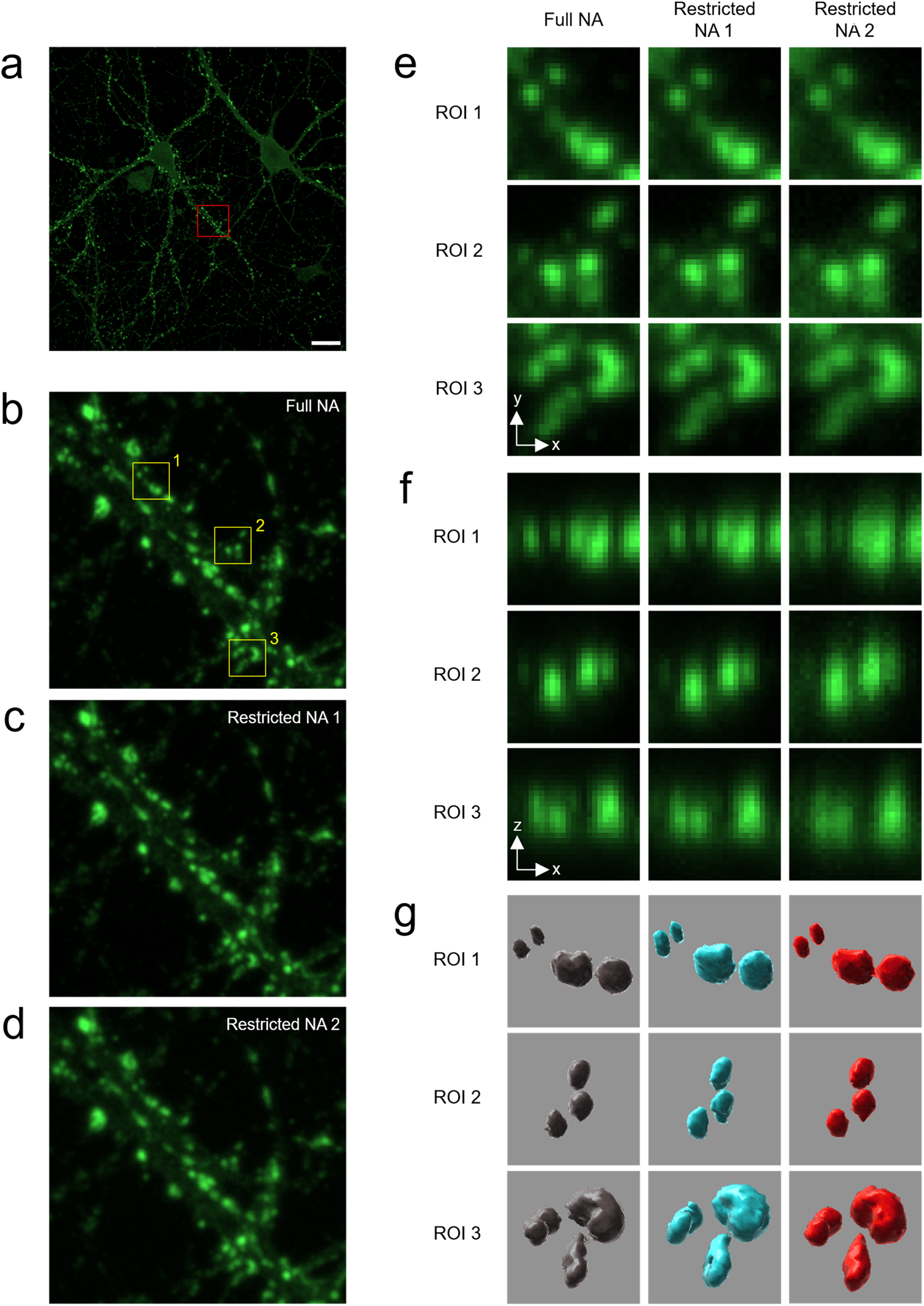
Changing dendritic spine puncta volumes of cultured neurons imaged under systematically reduced microscope resolution conditions. **a**, Full FOV Green channel maximum intensity Z-projection image of neurons with fluorescently tagged AZ vGLUT proteins acquired with Objective 1. Scale bar = 20 μm. **b-d**, Zoomed in views of a single dendritic process from the red boxed ROI in **a** for the Full NA, Restricted NA 1 and Restricted NA 2 emission pathway iris conditions respectively. **e**, Further digitally zoomed views of individual puncta from the yellow boxed ROIs in **b**. **f**, Side-view XZ intensity projections and **g**, 3D Imaris surface renderings of the same dendritic spine puncta from **e** under the various NA settings.

Alterations of the spine puncta shape and size were visually subtle and difficult to spot (again emphasizing the need for quantitative measurement and how our visual impressions of changing image features often *do not* correlate with quantitative variations in distance-based image measurements), but were discernible in the extreme case of Restricted NA 2, especially when examining zoomed-in, axial side views of individual puncta (Fig. 4f) or when spine puncta were surface rendered in 3D and presented side-by-side (Fig. 4g). Fourier-domain MTF analysis of these cellular images showed a decrease in the lateral cutoff frequency (Supplementary Fig. 2), mirroring the microsphere results (Supplementary Fig. 1) and confirming that stopping down the emission pathway iris also increased blur in the fixed sample images.

Comparing the same presynaptic puncta volume populations between the two extreme NA conditions (Full NA and Restricted NA 2) showed that both could be described by a log-normal distribution (fitted solid curves, Fig. 5a) with the majority of puncta having volumes in 0.150 to 0.300 fL range, consistent with previous reports imaging the same protein by cryo-electon tomography and super-resolution stimulated emission depletion (STED) microscopy^72,73^. When the iris was closed down to the Restricted NA 2 condition, we observed a measurable increase in puncta volumes, but mostly biased towards the smaller puncta volumes, as indicated by the shifted curve peak position between the two distributions. In conjunction with the model presented in the Supplementary Text, the more pronounced volume growth of smaller puncta makes sense: Resolution degradations mostly impact measurements of image features which are comparable in size to the resolution limit. Beyond this size, the blurring effect on the volumes of spherical objects having a diameter that exceed the resolution limit by ∼2-3x is negligible (see Equation S10 in Supplementary Text) and explains the distribution range of the tail regions of both histograms appearing similar (Fig. 5a). Both the modal puncta volume of the Full NA condition (Fig. 5b - 0.190 fL, vertical dashed line) and the peak distribution shift (19%, horizontal dashed line) almost intersect perfectly with the theoretical curve, and the theoretical curve itself passes roughly through the middle of the scattergram data points, thus demonstrating good agreement between theory and measurement for this control experiment.

**Fig 5.**
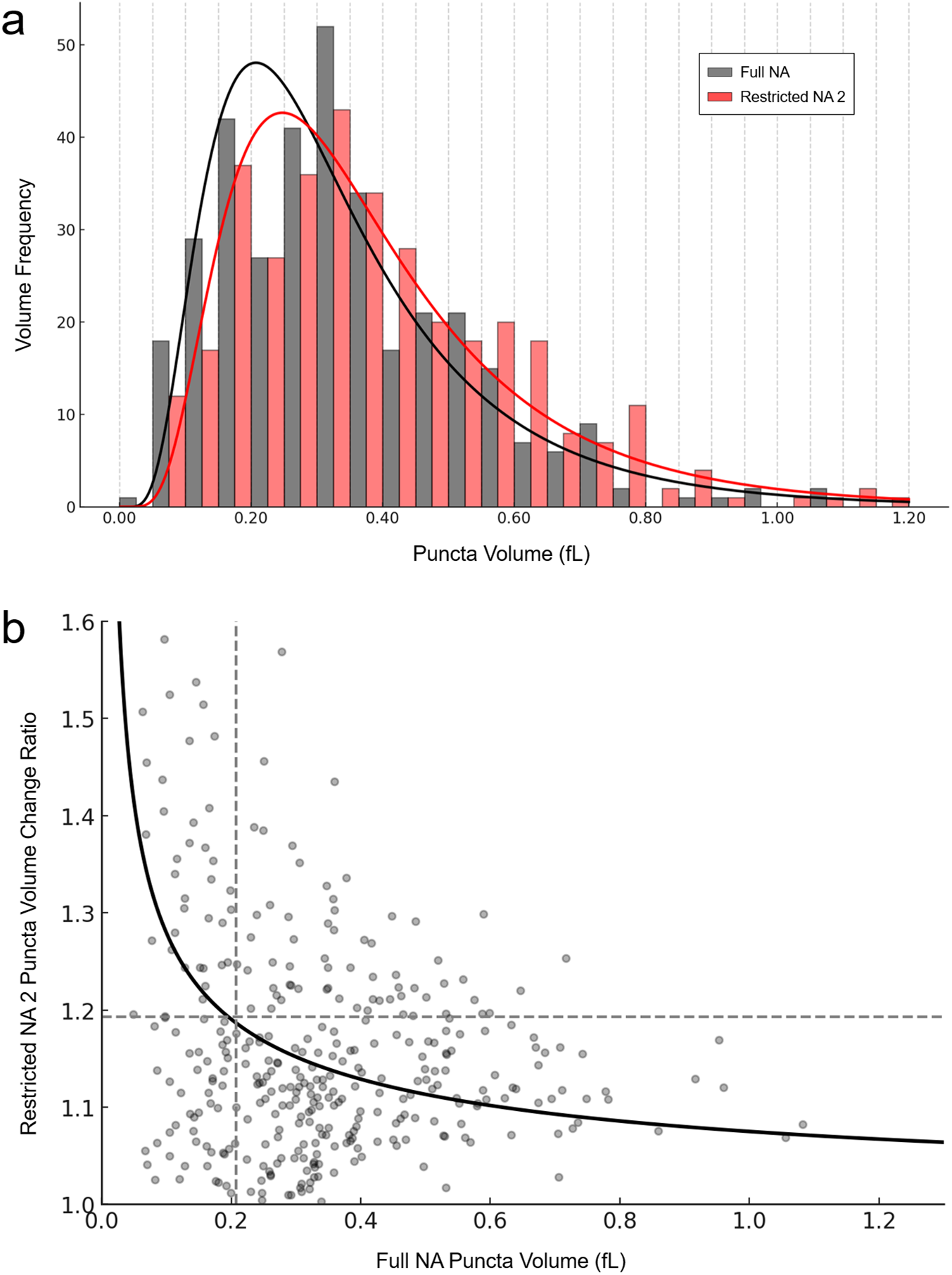
Dendritic spine puncta volume inflation for the resolution control experiment. **a**, Histograms of measured puncta volumes for the Full NA and Restricted NA 2 conditions extracted from the full FOV neuron images presented in Fig. 4. **b**, Scattergram plot of puncta volume change ratio vs. initial Full NA puncta volume for every identified dendritic spine punctum. The solid black curve is the theoretical curve associated with these resolution-induced volume changes (Equation S11, Supplementary Text) for different sized spherical objects. The vertical dashed line shows the measured peak (mode) puncta volume of the Full NA condition (0.207 fL) while the horizontal dashed line is the distribution peak shift from Full NA to Restricted NA 2 (19%).

To visualize the size-dependent puncta inflation depicted in Fig. 5 and predicted by Supplementary Equation S10, we simulated 3D images of fluorescent spheres convolved with 3D Gaussian ellipsoid-shaped PSFs of increasing width (reduced effective NA). The apparent simulated volumes follow the analytic trend where features whose diameters are comparable to the PSF extent expand markedly as blur increases, whereas objects larger than ∼0.5 µm are minimally affected (see Supplementary Fig. 3). This explains why the shift in Fig. 5a is concentrated among smaller puncta and why the volume-change ratio in Fig. 5b decays with initial size.

Our second resolution experiment replicated the imaging procedures of the control experiment. but utilizing 3 similar objective lenses mounted on the same microscope instead of directly manipulating the resolution of one objective lens. The three objectives were all Nikon 60× Plan Apochromats with an NA of 1.40 (Objective 2 and 3) or 1.42 (Objective 1, Fig. 6). As before, we used 200 nm sized fluorescent microspheres to characterize the PSF FWHM lateral and axial extents associated with each objective lens and found they all exhibited relatively spatial homogenous resolutions across the FOV, but with distinct and measurable differences across objectives, reflecting the natural variations in objective lens quality (Fig. 7). The average lateral resolutions of each objective lens were comparable and about 23-32% larger than their theoretical values based on their operation wavelength (529 nm) and NA (1.40 for Objective 2 and 3, 1.42 for Objective 1). Objective 2 showed slightly larger axial fits and visual inspection of the images showed an asymmetric “banana” shaped aberration in its PSF, possibly indicative of damaged or defective internal lens elements (Fig. 6a and Fig. 7a). Surprisingly, Objective 3 possessed the best (smallest) overall lateral and axial FWHM, with measured resolution data pairs clustering in a cloud closest to the 1-1 region of the FWHM scattergram (Fig. 7b and c), even though Objective 1 had a slightly higher NA and was the latest generation model of this lens (LambdaD vs. Lambda series). At the same time Objective 1’s lateral and axial resolution was more consistent across the full FOV (Fig. 7a). Interestingly, all three of these objective lenses would be considered “acceptable” in terms of resolution performance according to the QUAREP-LiMi recommendation standards.

**Fig 6.**
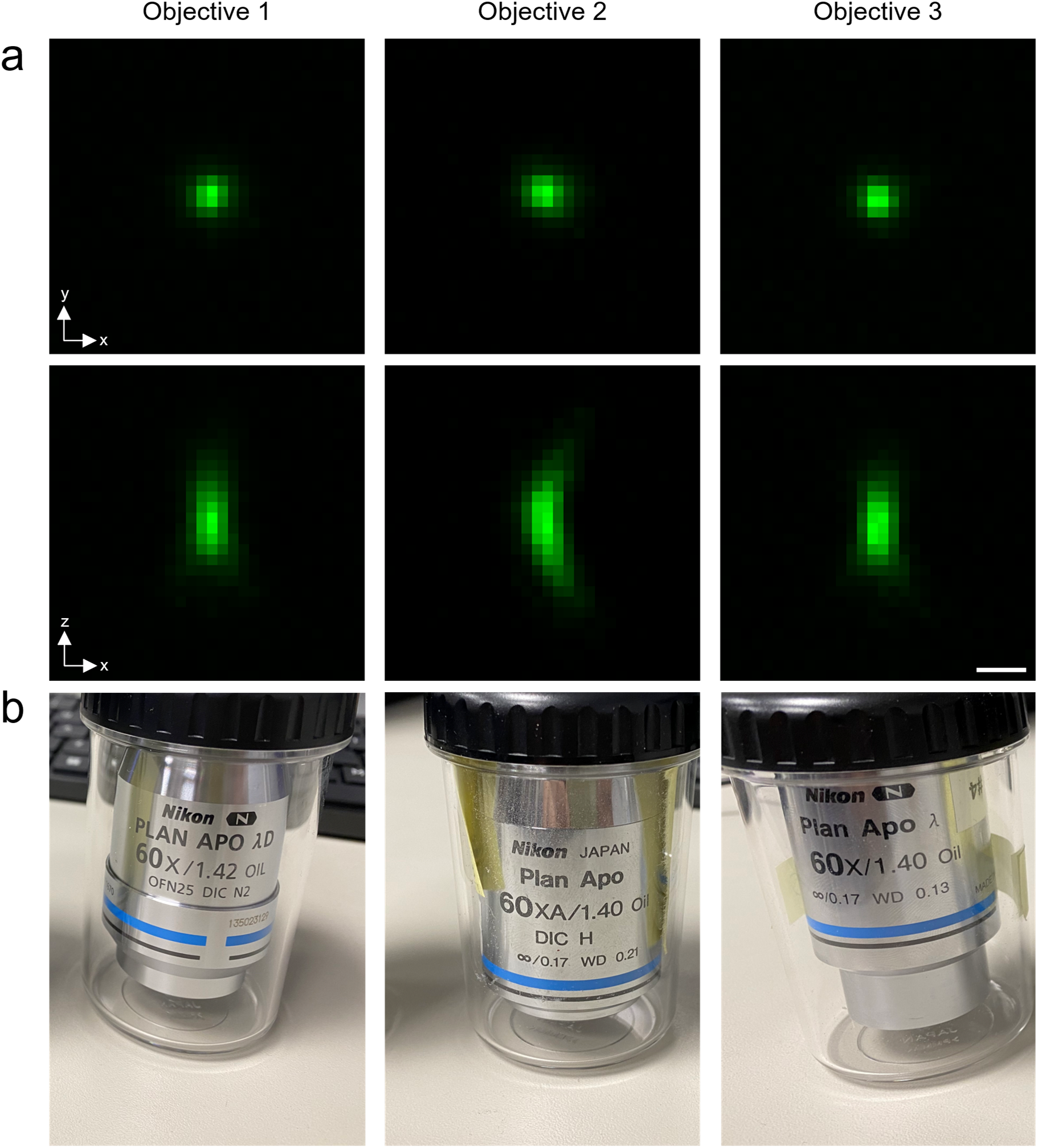
PSF comparison across three 60× oil immersion objective lenses imaging the exact same fluorescent microsphere. **a**, XY (top row) and XZ (bottom row) maximum intensity projections of the exact same 200 nm fluorescent microsphere imaged on the same microscope with three different 60× oil immersion objective lenses. Scale bar = 0.5 μm. **b**, Photographs of the objective lenses used for this study.

**Fig 7.**
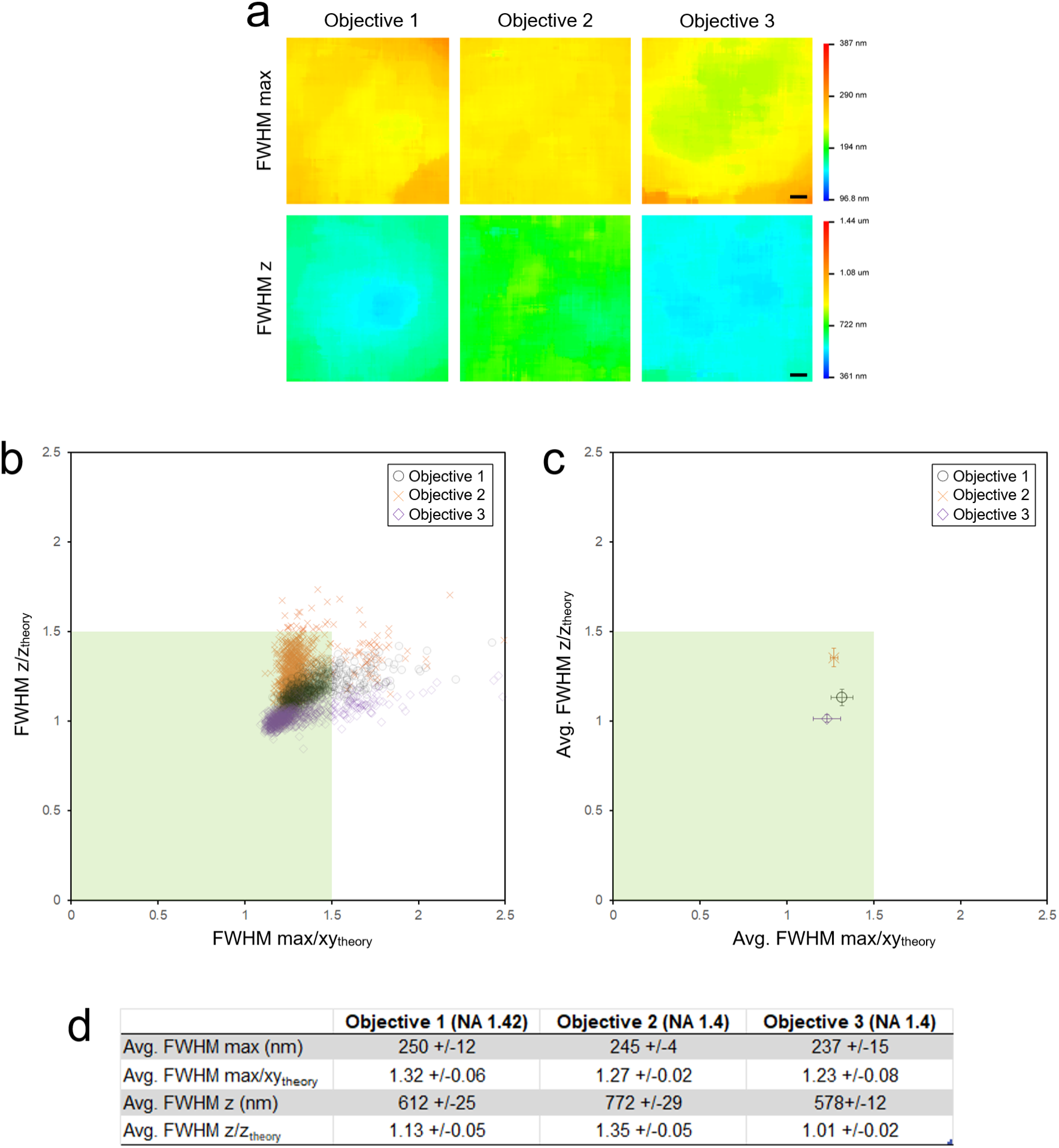
Full FOV lateral and axial resolution characterization of Objective lenses 1-3. **a**, PSFj lateral (FWHM_max_) and axial (FWHM_z_) resolution heatmaps of Objective 1, 2, and 3. Scale bars = 20 μm. **b**, For each objective, the lateral and axial FWHM values of each isolated microsphere from the FOV were normalized by their theoretical values and plotted against each other. The green box delimiting the 1.5x theoretical resolution ratios match the recommended microscope resolution performance tolerance limits of Faklaris et al.^12^ **c**, The centroids of each data “cloud” in **b** plotted the same way representing the average resolution performance across the entire FOV for each objective lens. **d**, Average FOV resolution FWHM measurements in tabular form.

Again, lateral Fourier domain MTF analysis of the microsphere maximum intensity Z-projection images validated these spatial domain PSF findings for the three objective lenses and showed smaller resolution differences than we obtained in the control experiment by restricting the NA of Objective 1 (Supplementary Fig. 4).

Given the subtle differences when measuring well-defined microspheres, we expected that the images of our biological specimen acquired with each objective lens would appear nearly identical, and indeed the example neuron images shown in Fig. 8 are nearly impossible to distinguish visually (although the banana-shaped PSF of Objective 2 does manifest itself more clearly in the orthogonal and 3D surface rendered views of the smallest presynaptic puncta (Fig. 8f and g).

**Fig 8.**
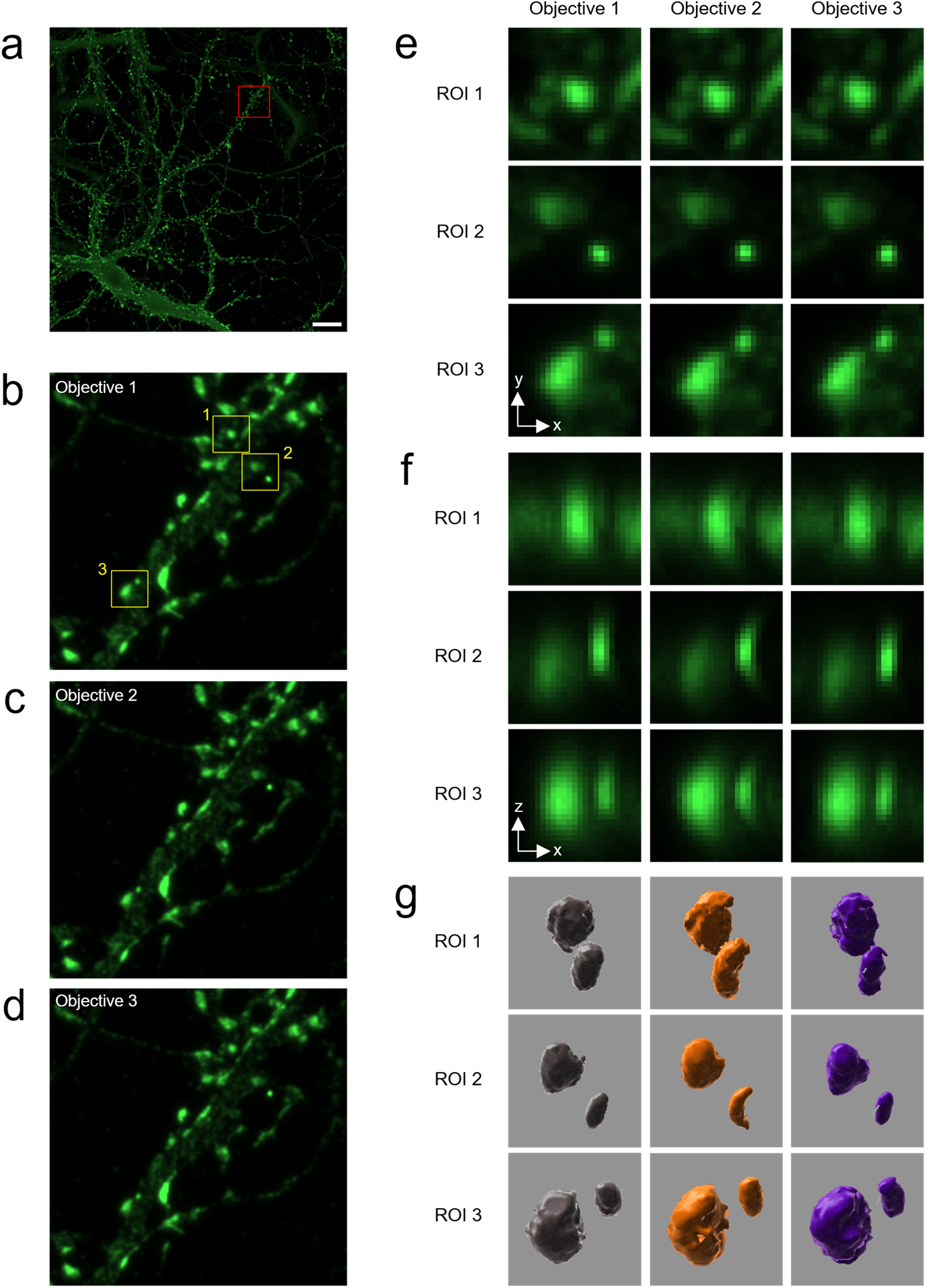
Comparative neuron imaging across Objectives 1-3. **a**, Whole FOV image of presynaptic immunolabelled vGLUT proteins in cultured hippocampal neurons from Objective 1. Scale bar = 20 μm. **b-d**, Zoom in view of red boxed ROI from panel A for all three objectives. **e-g**, Yellow boxed sub-ROIs 1-3 from panels **b-d** for each objective lens depicting individual vGLUT puncta shown laterally **e**, orthogonally **f**, and as surface rendered volumes **g**.

However, an equivalent statistical analysis of the measured puncta volumes observed in the images acquired with each objective lens reveals that the volume distribution differences are significant and correlate with the ranking of measured objective lens resolutions (Objective 3 < Objective 1 < Objective 2), as illustrated in Fig. 9. Plotting all measured puncta volumes for each objective, we were still able to observe measurable volume differences when comparing the same central FOV with significant differences in Mann-Whitney U tests between Objective 1 and 3 and Objective 2 and 3, but not between Objective 1 and 2 (Fig. 9c). The results of this second resolution experiment show again that apparent puncta volumes scale with the instrument’s PSF and motivate resolution-based instrument normalization of intra-channel image readouts (using measured PSF FWHM) to ensure that measured feature differences obtained across systems reflect biology rather than optical blur.

**Fig 9.**
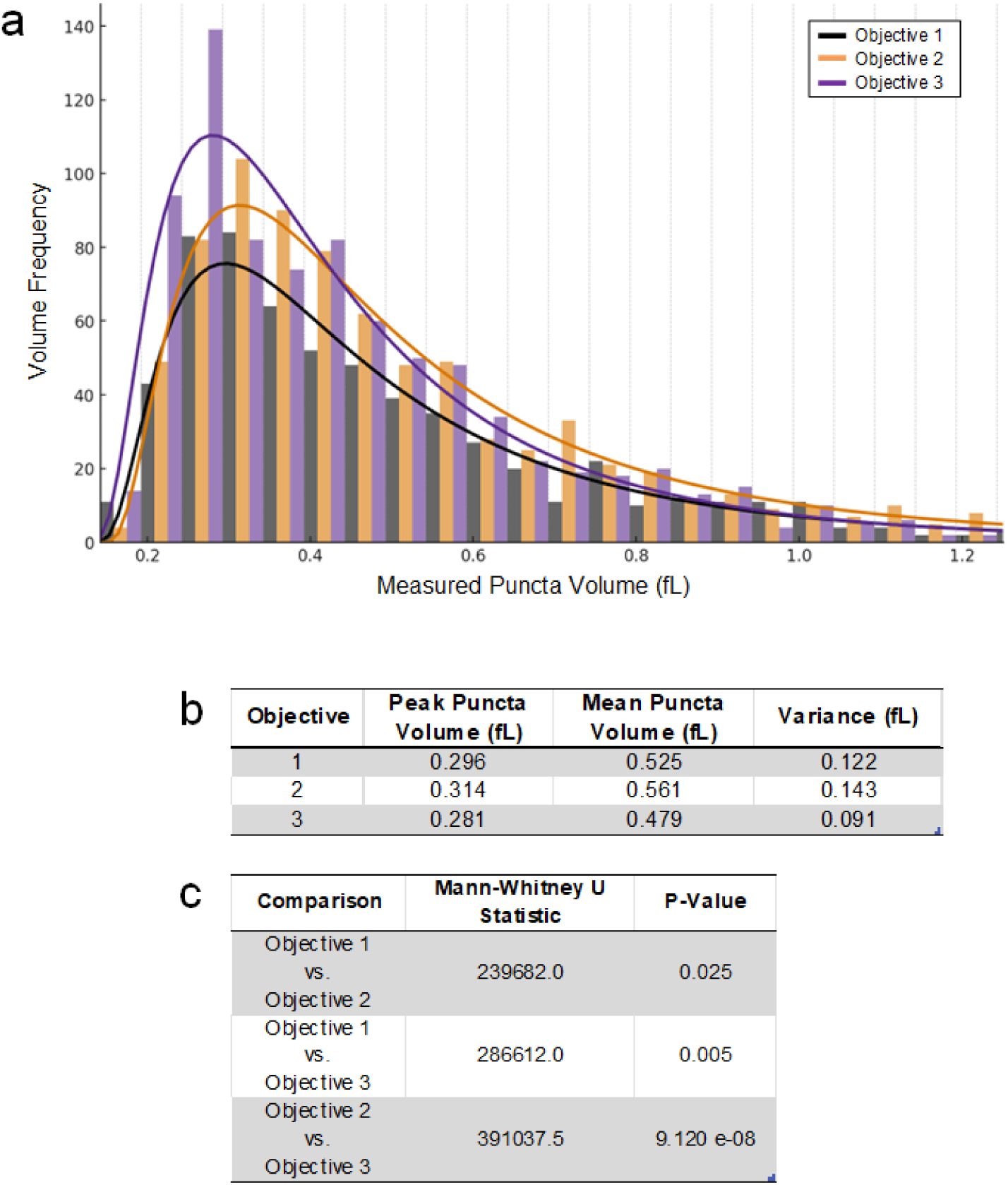
Objective-dependent differences in measured presynaptic vGLUT puncta volumes. **a**, Distributions of presynaptic puncta volumes extracted from the whole FOV shown in Fig. 8 across the 3 objectives. All three objectives showed similar distribution shapes. **b**, Statistical analysis of the modal (peak) and mean volumes and their variance for each objective. **c**, Mann-Whitney U pairwise tests of the three objective volume results, demonstrating that the measured differences are statistically significant in two of the three pairwise cases.

### How channel co-registration impacts sub-cellular inter-channel image measurements of colocalization and volume overlap

To isolate the optical contribution to two-channel image measurement readouts, we first established where each objective lens was best registered and how much the channel mis- registration varied across the detector FOV. Full FOV chromatic aberration maps derived from two-color microsphere QC calibration tests showed that lateral chromatic shifts dominate and grow toward the periphery, while axial shifts for these three objective lenses were smaller and more uniform (Fig. 10a). These lateral/axial chromatic aberrations are unified into a single *scalar* Chromatic Displacement Index (CDI) ratio, x – a channel pair co-registration QC metric that combines lateral and axial offsets after normalizing by the measured PSF widths (FWHM), so that χ reflects mis-registration in units of the system’s actual resolving power (see Supplementary Text and Equation S13). This is in contrast to the original metric definition recommended by Faklaris *et al*. with the co-registration ratio applying the *theoretical* lateral and axial resolutions for normalization instead in an instrument-agnostic manner^12^. Switching the CDI normalization to the *measured* PSF FWHM (per-objective and per-FOV location) advantageously makes x an instrument- and site-specific detection accurate QC metric. It displays whether the mis-registration is large relative to what the microscope actually resolved in that image dataset. This adjustment is also consistent with the quantitative framework presented in the Supplementary Text, which shows that two-channel colocalization/overlap readouts depend exponentially on inter-channel mis-registration using PSF-normalized units. A broader PSF lessens apparent registration penalties, whereas a sharper PSF amplifies them.

**Fig 10.**
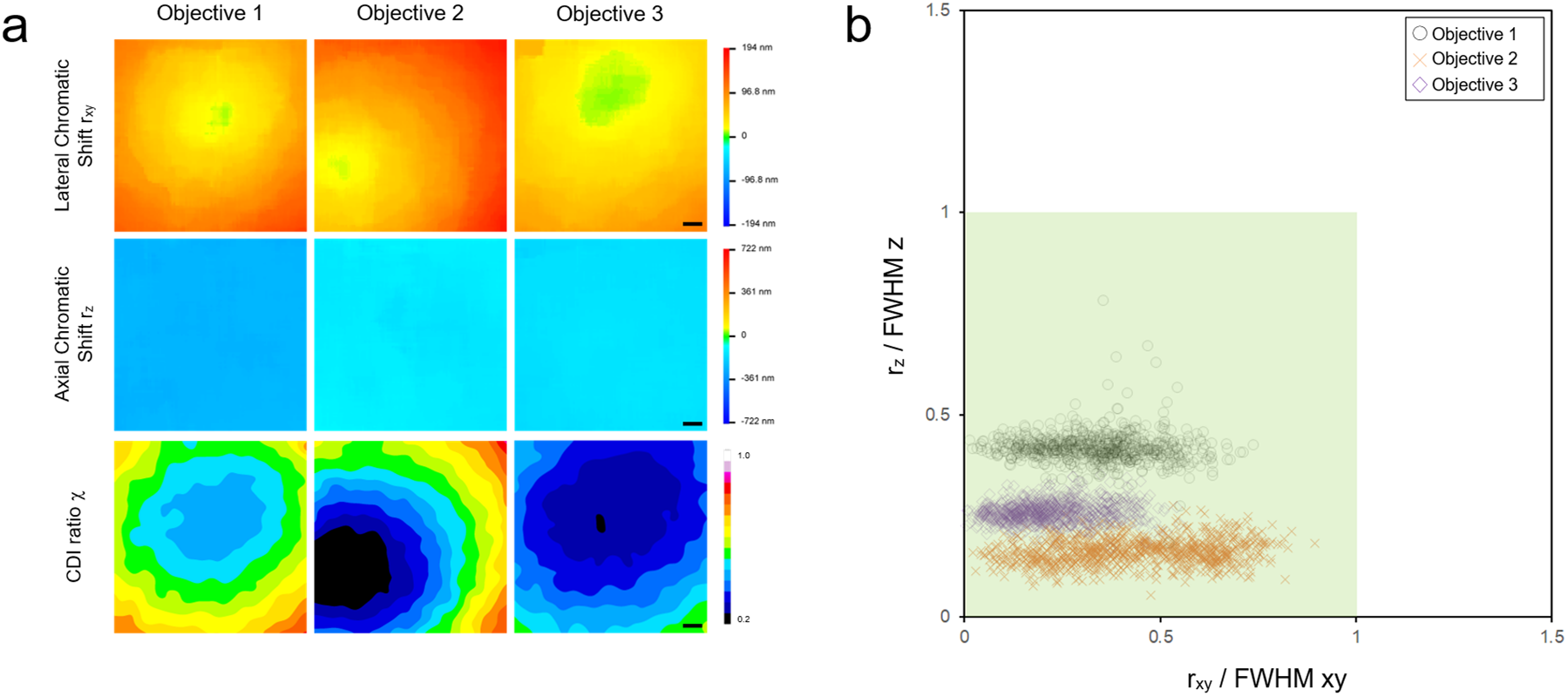
Full field of view objective lens chromatic co-registration characterization with measured PSF normalization. **a**, Green-Yellow channel pair heatmaps derived from two-color microsphere QC calibrations for objective lenses 1-3. Top row: Absolute lateral chromatic displacement magnitude *r_xy_* (nm) map. Middle row: Absolute axial chromatic displacement magnitude *r_z_* (nm) map. Bottom row: Scalar CDI ratio χ (unitless) map, which combines lateral and axial shifts after normalization by the corresponding measured PSF FWHM values for the respective objective lens. The CDI ratio heatmaps use a different 16-step discretized color look-up-table (LUT) spanning a range of 0.2-1.0; Lower χ values (black < blue < green … < white) indicate better inter-channel registration. Scale bars = 20 μm. Across objectives, lateral offsets dominate and increase radially toward the periphery, whereas axial offsets are small and comparatively uniform. Objective 3 exhibits the lowest χ over the largest fraction of the FOV (extensive dark blue contoured zone, indicating superior co-registration performance; Objective 2 shows a pronounced region of near-perfect co-registration (black zone) that is significantly offset from the FOV center; Objective 1 is radially symmetric like Objective 3 with χ values still within the recommended tolerance over the entire FOV. These heatmaps enable users to select both the objective and the specific FOV sub-region that meets the precision required for a given experiment. **b,** Scatterplot of the axial (*r_z_*) vs. lateral (*r_xy_*) channel displacements of every measured microsphere within the FOV normalized by the respective objective lens’ measured lateral and axial PSF FWHM (FWHM_z_ and FWHM_xy_). The shaded green box denotes the acceptance window (as recommended by Faklaris et al.^12^) where both normalized shifts are < 1. Microsphere channel displacements from Objective 3 cluster closest to the origin, consistent with panel **a,** identifying this objective as the best choice for colocalization studies among the three optics, while Objectives 1 and 2 remain usable since their data clouds are well within the tolerance zone as well.

Thus, measured PSF normalization aligns the metric with the true sensitivity of the acquisition rather than an ideal theoretical limit that the system may not reach (e.g., due to aberrations or depth-dependent blur), and it avoids over- or under-penalizing the same geometric shift across different instruments or ROIs. For continuity with prior practice, we also report the theory- normalized CDI co-registration ratio (x_theo_) spatial maps (using the shorter-wavelength channel theoretical FWHMs) in Supplementary Fig. 5. Theoretical resolution normalization values facilitate instrument/optical component benchmarking, whereas the measured-PSF normalization we adopt for modeling is a more appropriate scaling for cross-study comparisons, as well as estimating experimental detectability and colocalization bias in the image data actually acquired. Objective 3 maintained low x over the largest area making it the best overall objective lens for inter-channel overlap image measurements, Objective 1 was intermediate, and Objective 2 exhibited a far off-center “sweet spot” of near-perfect registration with poorer regions elsewhere. These co-registration heatmap trends and observations were corroborated by the similarly measured PSF FWHM-normalized axial (*r_z_*) vs. lateral (*r_xy_*) channel displacement scatterplot (Fig. 10b). This plot again suggested the superiority of Objective 3 for colocalization studies since the centroid of the data cloud associated with this objective lens lay closest to the origin (which represents the state of perfect channel pair registration). In contrast, the centroids of the data clouds associated with Objective 1 and 2 were about equidistant from the scatterplot origin and indicated their differences in lateral versus axial co-registration performance. While such scatterplots are useful for rapid objective lens comparisons, they lack the spatial context that the CDI heatmaps provide, which is critical for choosing inter-channel analysis regions.

Guided by these co-registration QC heatmaps, we then acquired matched biological data from the same fixed pre- and postsynaptic immunolabeled neuron at two positions: the FOV center (ROI A1) and, after a motorized XY translation, the upper-left corner (ROI A2), repeating this center/corner confocal Z-stack pair acquisition for all three objectives under identical settings (Fig. 11). This experiment design held the biology constant while intentionally sampling different local channel registration states known from the QC CDI x heatmaps. Visually, the merged vGLUT-Homer overlays suggested more overlap/colocalization at the image center than at the corner (Fig. 12), consistent with the expectation that registration errors can modulate the apparent colocalization or volume overlap.

**Fig. 11.**
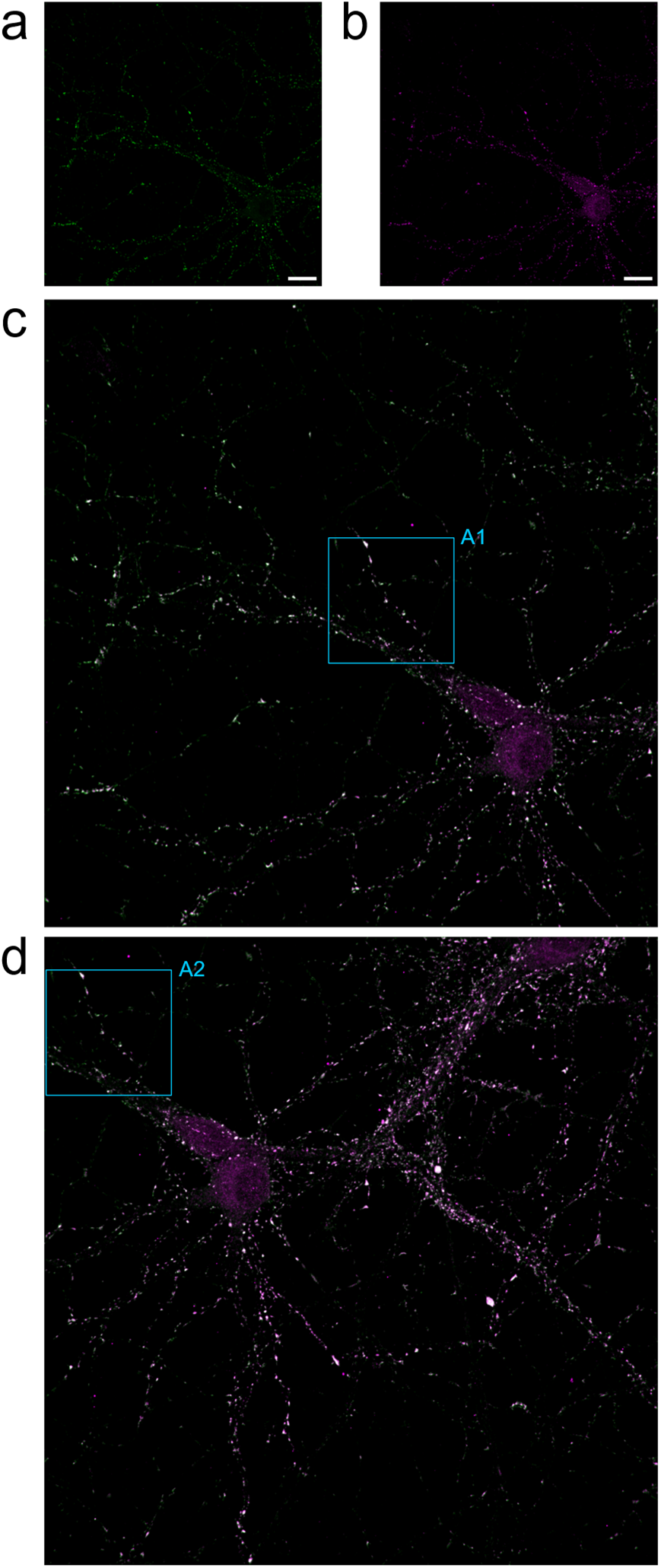
Central vs. edge two-color neuron images and ROI selection for colocalization analysis under spatially varying chromatic co-registration. Single-channel maximum-intensity Z-projections from 3D confocal stacks of fixed neurons immunolabeled for vGLUT (**a**, Green channel, green LUT) and Homer (**b**, Yellow channel, magenta LUT), acquired with Objective 1. Scale bars = 20 µm. **c**, Color-merged overlay of panels **a**–**b** with cyan box labeled A1 marking the ROI used for quantitative colocalization analysis. **d**, The same neuron re-imaged after a motorized XY-stage translation that moved the neuron into the upper-left corner of the FOV. Both channels were re-acquired with identical settings. Cyan box labeled A2 denotes the ROI that was spatially identical to A1 in panel **c**. These ROIs were chosen to sample zones with different local chromatic co-registration quality (low-χ center vs higher-χ edge) as determined by QC-derived CDI co-registration heatmaps (see Fig. 10).

**Fig 12.**
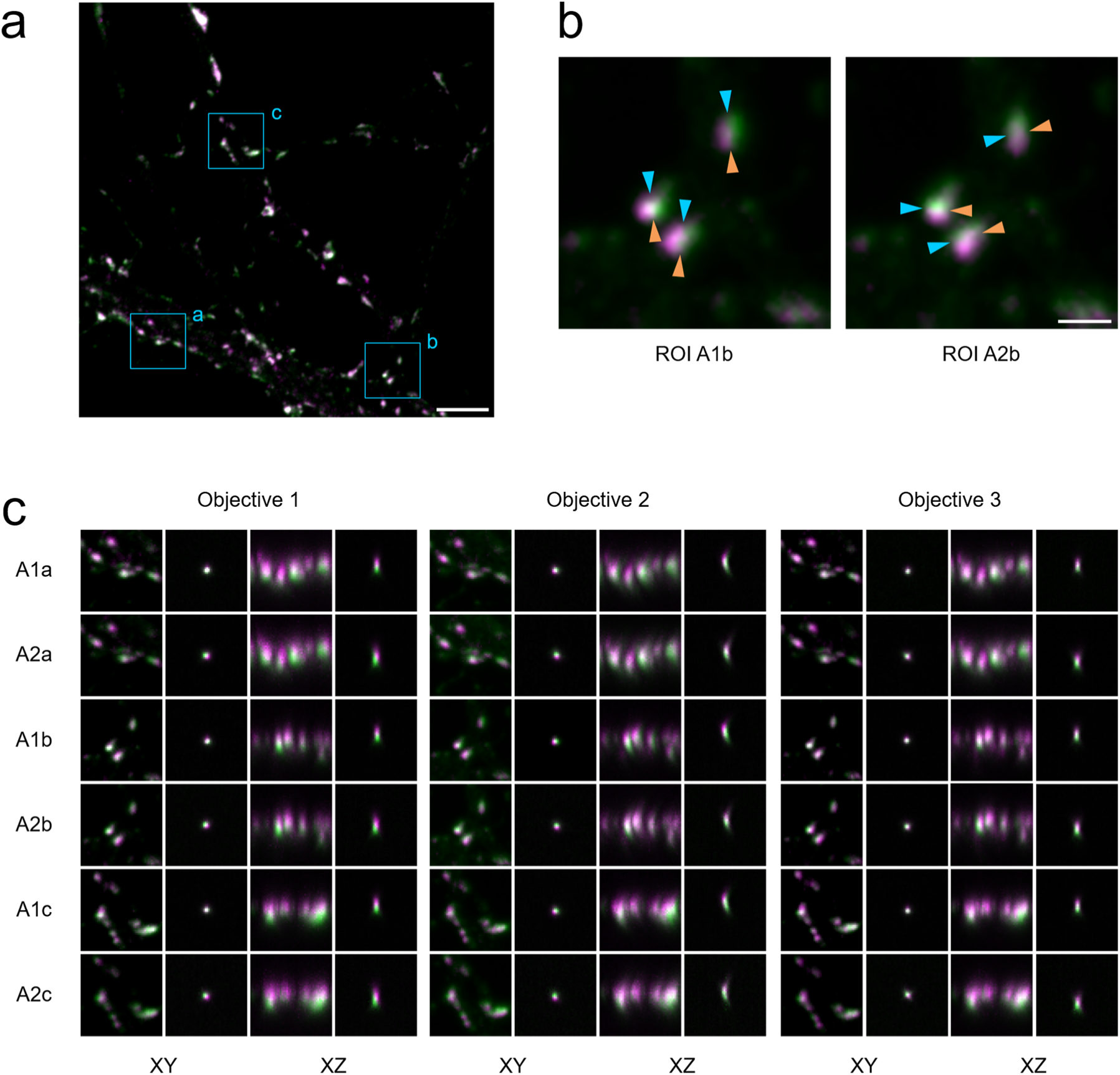
Visual effects of objective- and position-dependent co-registration on synaptic puncta overlap and apparent cleft orientation, with matched bead-QC orthogonal views (vGLUT, green LUT; Homer, magenta LUT; overlap appears white). **a**, Two-color confocal maximum intensity Z-projection overlay acquired with Objective 1; cyan boxes (a–c) mark sub-ROIs chosen for orthogonal inspection from ROI A shown in Fig. 11. Scale bar = 5 μm **b**, Close-ups of sub-ROI b from the center (A1b, left) and upper-left corner (A2b, right) of the FOV acquired with Objective 2. Cyan/orange arrowheads indicate the apparent pre- versus postsynaptic sides to show that the cleft orientation can change between positions, which is a consequence of differences in the local co-registration vector (direction/sign). Subtle differences in puncta overlap (white) are also evident with each objective lens and position. Scale bar = 1 μm **c**, For each sub-ROI pair (rows A1a/A2a, A1b/A2b, A1c/A2c), XY and XZ views of the synaptic puncta are shown for Objectives 1–3, alongside matched microsphere-QC XY/XZ profiles taken from the same FOV locations in the calibration data. Center ROIs generally show tighter apposition (more white) and more symmetric XZ profiles, whereas corner ROIs show increased channel separation and orientation bias that mirror the local QC measurements (see Fig. 10a).

Identical image pairs of the exact same neuron at the same two FOV positions as shown for Objective 1 in Fig. 12 were also acquired with Objectives 2 and 3. These paired ROIs provide a controlled test of how spatially varying co-registration affects vGLUT–Homer colocalization readouts. Orthogonal views (as depicted in Fig. 12c) made these microscope channel registration dependencies explicit at the single dendrite punctum scale. Sub-ROIs drawn from A1 (center) and A2 (corner) revealed (i) visible changes in punctum overlap and, in some cases, (ii) changes in apparent cleft orientation for the same pre/post punctum pair, emphasizing that these effects were purely instrument-driven. For Objective 2 in particular, ROI center-to-corner comparisons showed an orientation change of the vGLUT↔Homer vector and a reduced colocalization/overlap. The matched microsphere QC views taken from the same FOV locations with this objective exhibited the same axial “banana-shaped” XZ signature, directly linking the biological appearance of the punctum to the objective’s aberration behavior, as was presented in Fig. 6a and Fig. 8.

These emergent biases of vGLUT-Homer protein complexes under the varying co-registration imaging conditions align with the analytical framework developed in the Supplementary Text. For covariance-like (e.g., Pearson’s correlation coefficient, PCC) and geometric inter-channel overlap type image measurements (ie: a typical 2-channel synaptogenesis assay), channel mis- registration enters as a multiplicative exponential attenuation of the idealized “no-shift” or zero channel shift value. The model predicts two robust behaviors: (i) For fixed object sizes, worse co-registration (larger channel displacements offsets) decreases intensity product-based (Hadamard) image measurements (PCC, volume overlap, etc.^74^), and (ii) measurement sensitivity depends on object-to-PSF size via a size comparison factor, α. Resolution blur (larger effective width, larger α) reduces the impact of a given channel shift, which is why an imaging system with more resolution blur can *appear* to have higher colocalization for the same underlying biology and registration error.

In Fig. 13, we quantify this experiment to model link by plotting the measured *change* in colocalization between ROIs A1 and A2 against the measured change in local co-registration for each objective lens, using Equation S23 (derived in the Supplementary Text). For each punctum we computed a Dice symmetric volume-overlap colocalization score^75,76^ and compared the edge-to-center colocalization ratio to the change in co-registration (Δχ²) between ROIs A2 and A1. We then overlaid theoretical predictions from our Gaussian overlap model using object-to- PSF size ratios (α) derived from the measured PSF and puncta volumes. The analysis indicated that across objectives the A1→A2 colocalization differences follow the trend extrapolated from Equation S23 within error bars for all three lenses, consistent with the monotonic, size-modulated attenuation predicted by the model. Reporting results with measured PSF normalization (Fig. 13a) yielded the best alignment between theory and data and most faithfully reflected instrument performance.

**Fig 13.**
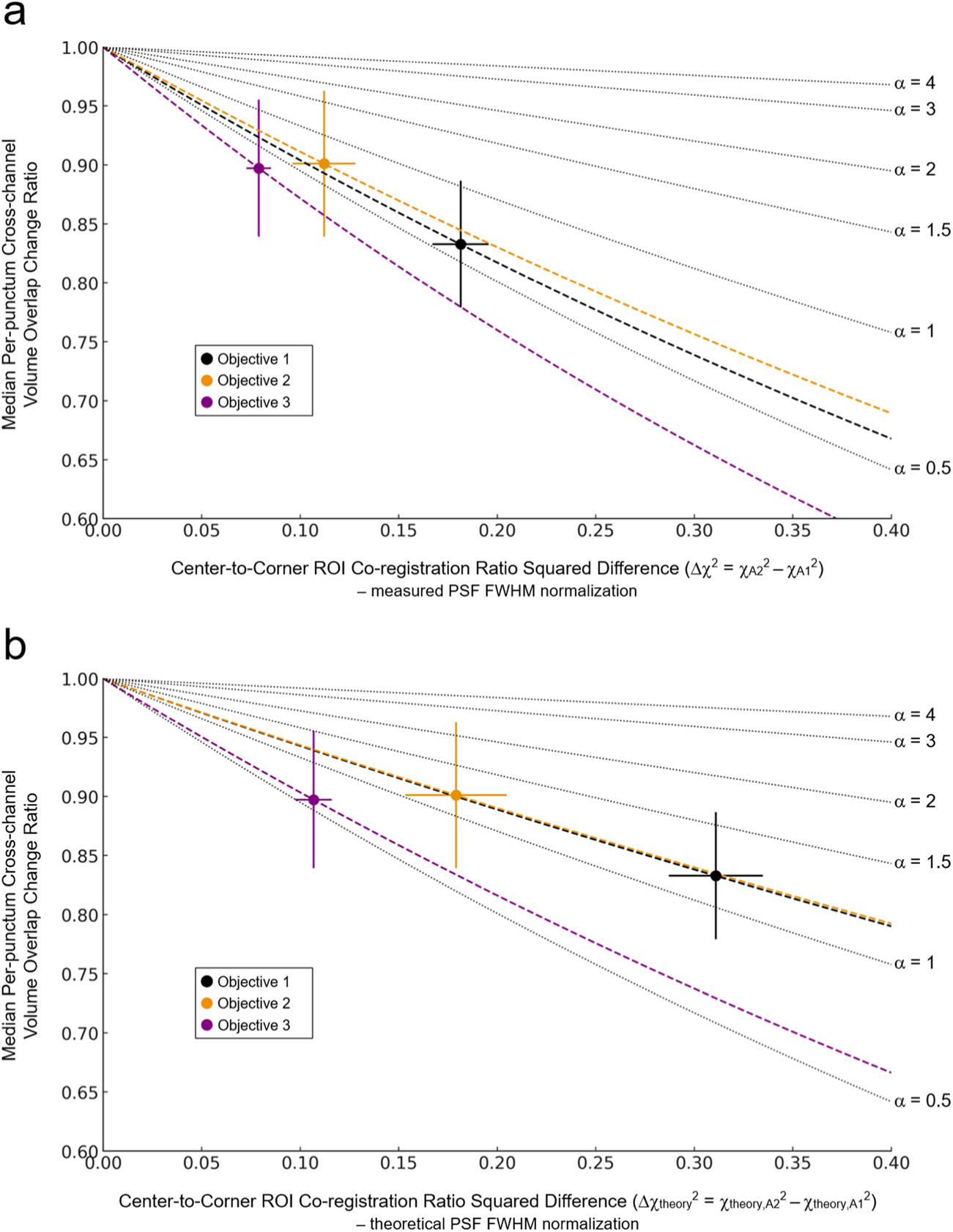
Inter-channel registration error predicts loss of puncta colocalization overlap. **a**, Dice (*C*) volume overlap coefficient ratio (*C*_A2_/*C*_A1_) versus change in co-registration error Δ(χ^2^)=χ_A2_^2^−χ_A1_^2^ with measured PSF FWHM normalization. Here, *C* denotes the Sørensen–Dice overlap coefficient between the 3D binary masks of the green and yellow channel punctum for each matched object:

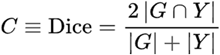 For each objective, 15 paired vGLUT-Homer puncta (A1↔A2; total N=45) visible in both channels were analyzed. Data points show the median of the per-punctum ratios and vertical uncertainty bars are bootstrap 95% confidence intervals for the median, while horizontal uncertainty bars are standard errors (SE) for Δ(χ^2^) propagated from the ROI-level uncertainties. Scalar co-registration χ values were computed as the mean +/− standard deviation (SD) of pixel values in the co-registration heatmaps of Fig. 10a, restricted to the same A1/A2 ROIs used in Fig. 12c–d. These ROI means (and SDs) feed Δ(χ^2^) and its SE. Dotted black curves show the Equation S23 reference family curves for different object to PSF size ratios for α={0.5,1,1.5,2,3,4}. Dashed colored curves are per-objective fits of α. **b**, The same analysis as in panel **a**, but with theoretical PSF FWHM normalization. Replacing measured PSFs with theoretical values shifts the points rightward (larger Δ(χ^2^), and the fitted α values correspondingly increase (two lenses move to α∼1.1−1.2; one remains <1).

Together, Fig. 10 - Fig. 12 establish the instrumental origin of the observed changes in inter- channel puncta overlap and orientation for the same neuron, and Fig. 13 provides the quantitative, model-consistent calibration that ties local co-registration to the magnitude of colocalization loss - another demonstration of how a critical microscope QC metric couples to and influences a real sample image measurement. As was noted with intra-channel distance- based measurements (e.g. volume), the QC metric matters most when the cross-channel feature measurement is comparable in size to the QC metric value (lower α).

## Discussion

A central aim of this study was to demonstrate directly and unambiguously that routine light-microscope QC metrics are not merely ancillary indicators of instrument “health” or guardrails for data consistency, but also determinants of quantitative image readouts. Since QC metrics set the scales at which features are sampled, changes in a QC value propagate predictably into the reported measurement with the direction and magnitude of the bias governed by how the feature compares to the metric (e.g., object size relative to the PSF; inter-channel displacement relative to object extent/overlap). This transformation of the data is bidirectional: biological inferences drawn from an image readout are valid and interpretable *only to the extent that the relevant QC metric remains stable over the measurement interval*. We showed this link concretely for two archetypal metric–measurement pairs: Image resolution and intra-channel, distance-based morphometrics (e.g., volume) and inter-channel co-registration and covariance-like and overlap-based indicators of colocalization.

In our constricted objective NA, resolution-reducing control experiment, the apparent presynaptic puncta volumes intuitively increased as the resolution of the system decreased. A more important question is if these correlations could have been predicted based on the observed changes in the QC samples alone. A 3D Gaussian convolution model of the microscope PSF which depends on the variance-addition properties of this function is presented in the Supplementary Text and explains how intra-channel distance-based image measurements (feature volume in this case) of punctate dendritic spines *intrinsically depend* on the system resolution, *but in a size-dependent manner*. Practically, even a minor broadening of the PSF, or a small optical misalignment that widens the effective blur made these objects appear >30% larger than the Full NA, well-aligned conditions, even for structures larger than the diffraction limit. Such inflation is easy to miss visually but can flip or confound experimental conclusions if the instrument state drifts between conditions. The three 60× objective lenses we tested differed only subtly and all of them would have passed common resolution tolerance cut- offs (e.g., FWHM-based thresholds recommended in recent community guidelines), underscoring that “within tolerance” does not mean “measurement-equivalent”. Routine reporting of resolution via a simple FWHM from sub-diffraction sized fluorescent microspheres provides the minimal calibration needed to interpret and compare volume measurements across sessions and instruments.

We note that Objective 2, despite its clear inherent aberration, could be perfectly acceptable for certain imaging scenarios and depending on the nature of the experiment. As we showed in the control experiment, the measurement of volume is affected by characteristics of the PSF only when the specimen features under study are similar in size/separation distance to the PSF itself. In contrast, when the relevant structures under study are several PSF widths across (or separated by many PSF widths), the intensity signal is dominated by broad, slowly varying features that are essentially unchanged by modest blur, and accordingly, differences in resolution or mild aberrations have negligible impact on the readout (assuming adequate digital sampling and signal to noise ratio).

The same normalization approach can be applied to any intra-channel, distance-based image measurement and is not limited to neuronal cell structures. Many cell-biological readouts from fluorescence microscopy like organelle morphometry (length, area, volume, cluster size, and other spatial statistics such as nearest-neighbor/centroid separations) of mitochondria, lysosomes, endosomes, focal adhesions, clathrin-coated pit sizes, nuclear foci/condensates, etc. depend on fluorescence segmentation and are therefore susceptible to PSF-induced size inflation and sampling biases. A recommended and lightweight best practice then is to pair PSF calibration with the same variance-addition modelling used here to compute lens- and session-specific correction factors. This “instrument bias normalization” improves accuracy without reprocessing images and offers an alternative to full deconvolution (which can be time-consuming, expensive, and assumption-laden) while simultaneously avoiding over-sharpening artefacts and deconvolution pipeline dependencies. Because the model is explicit in PSF units, one can also re-project a measured volume/area/length/etc. into the PSF of another system or time point provided steps are taken to determine the object/feature to PSF size ratio (α) by a resolution/co-registration perturbation through some auxiliary experiment (like we have demonstrated here), or if α is known/estimated through some other empirical means (electron microscopy or super-resolution imaging for example). This allows a researcher to ask the question, “What would I have measured on a microscope with PSF X instead of Y?” This re- projection of data into the characteristics of another microscope enables cross-system comparability and corrects the very biases that would otherwise confound quantitative fluorescence measurements, without altering the underlying images. Indeed, in our second set of resolution experiments where the exact same fluorescent neuronal synapses were imaged with three different objective lenses with all other things being equal, we measured three distinct puncta distributions, again stressing that the measurement only makes sense and converges to agreement when cited along with or corrected by the measured system resolution characteristics according to the instrument bias normalization approach outlined in the Supplementary Text.

As a theoretical illustration of the proposed normalization practice, suppose Lab 1 images vGLUT puncta with a 60×, 1.42 NA objective lens and Lab 2 uses a 100×, 1.30 NA objective. Both labs segment volumes from the raw fluorescence. Reported medians differ by ∼15%, leaving reviewers to juggle two volume lists and two PSF tables to guess whether the data gap is biological or optical in origin. Re-projecting both datasets to a common reference PSF (e.g., the measured PSF from Lab 1, or a community “standard PSF”) removes the instrument term analytically using the closed-form inflation factors in the Supplementary Text (variance-addition formulas for length/area/volume under isotropic or anisotropic PSFs). If, after re-projection, the two volume distributions still exhibit a residual difference, that difference can confidently be identified as biological in origin (within the sampling error) rather than instrumental. Crucially, this is achieved without altering images: The correction applies to the numbers using each dataset’s measured FWHM_xy,z_, and the results become directly comparable across experimental setups, time, and independent labs. By coupling every quantitative intra-channel image measurement to its intrinsically related microscope QC metrics, the reporting of imaging results then becomes explicit and FAIR (findable, accessible, interoperable, and reusable)^77,78^, enabling reviewers and downstream users to compare effects across microscopes.

Across our dual-channel datasets we observed that the exact same pre- and postsynaptic punctum, imaged with different objective lenses or at different FOV locations, could show measurable changes in apparent pre→post volume overlap. These shifts co-varied with the QC scalar maps of local chromatic co-registration χ. In this purposefully arranged experiment, we also confirmed that modest, spatially varying offsets could reorient the AZ↔PSD vector without any biological change, rendering both the true cleft orientation and protein colocalization ambiguous in the absence of related QC information, as well as potentially causing biased downstream interpretations of this synaptogenesis assay. Again, the approach to disentangle inter-channel instrumental dependencies from real sample features is not limited to neuronal cells. Applying Δχ-derived corrections (or constraining analysis to low-∣Δχ∣ regions) stabilizes orientation and overlap image readouts across objectives and positions. This behavior matches the 2-channel Gaussian PSF overlap model presented in the Supplementary Text (Equations S20 and S23). In PSF-normalized units, mis-registration enters as a predictable, multiplicative exponential on the idealized no-shift inter-channel measurement,

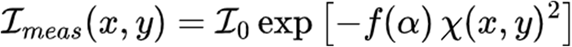

Thus, once χ is mapped and the object-to-PSF scale α is known or reasonably estimated via complementary experimental techniques, one can *a priori* forecast precision loss for any overlap/colocalization readout. As an illustrative back-of-the-envelope estimate, a ∼150 nm chromatic shift dampens the overlap of diffraction-limited synaptic puncta by ≈40%, but by only ≈6% for structures ∼3× larger than the PSF core. In this sense, QC co-registration heatmaps act like transfer functions that guide ROI selection and objective choice, making cross-system experimental comparisons interpretable when Δχ and α are reported.

A striking implication of the model is that a microscope with broader PSF (due to aberrations or imperfect alignment) can show higher apparent two-channel overlap/colocalization for the *same* underlying registration error and biology. That is not a virtue of the instrument, but rather a manifestation of the α term in the equation above. The same Δχ^2^ produces a smaller mis- registration penalty when the effective width of the feature is larger relative to the PSF (larger α). Intuitively, a system experiencing blur due to degraded resolution acts as a low-pass filter, acting to smear intensity variations such that a fixed physical mis-registration is less visible/quantifiable to overlap/covariance-type measurements. Therefore, without accounting for PSF and object size, cross-system or cross-day comparisons can be biased and confounding.

The remedy is not to accept blur, but to measure it and either standardize it across datasets or explicitly correct for its effect using the model. Although most microscopists conducting research at the cellular level likely understand this effect, its quantitative connection to the microscope’s co-registration QC metric which we have demonstrated here is under-appreciated.

Our analytic links between QC and measurement rest on two simplifications. First, we treat both the PSF and the effective object as Gaussian distributions that are separable across axes. In most day-to-day conditions this is a useful and reasonable approximation since optical aberrations primarily alter the pre-exponential factor without changing the exponential dependence on Δ(χ^2^). However, strong PSF asymmetries, field tilt, or pronounced axial–lateral coupling would warrant a more complex axis-resolved treatment (for example by replacing the scalar form with lateral/axial terms, e.g., f(α_xy_), f(α_z_), and use a vectorized registration field rather than a single scalar χ value). Second, we analyzed ROIs where the biological offset between channels, δ, averages to ∼0 (e.g., pre/post markers centered across the cleft).. When δ≠0 because of orientated structures, the observed overlap/colocalization reflects the combined separation ∣**δ**±**Δr**∣. In those cases and depending on the geometry, microscope channel mis- registration can mask true offsets, or amplify them. The model framework generalizes by adding **δ** to the mis-registration term and, for mixed populations, modeling the distribution of **δ**. In practice, reporting lateral and axial PSF extents, the vectorized co-registration map, and any known biological offsets alongside α and Δ(χ^2^) keeps interpretations of these more complex imaging scenarios rational.

## Conclusion

Our study reframes routine microscope QC as determinants of quantitative readouts rather than just auxiliary health checks of an instrument. We derived analytical links from measured PSF blur and chromatic mis-registration to the bias they induce in common image measurements (volume, overlap/colocalization). We also showed that colocalization signals depend exponentially on inter-channel mis-registration in PSF-normalized units, and that normalizing by the measured PSF instead of the theoretical limit is both necessary and sufficient to bring data and theory into agreement across objectives and FOV positions. We described an instrument- normalization / re-projection workflow that removes the instrument contributions to these measurement errors, allowing one to ask, *“How would this dataset change and appear on a microscope with PSF and co-registration X instead of Y?”* without altering or transforming any of the acquired images. Together these elements convert QC from a passive checklist of instrument health into a predictive design tool for setting tolerances, choosing acquisition/analysis ROIs, and making cross-system comparisons that are genuinely biological.

Practically, the recommended application of QC metrics into an experimental workflow is simple. First measure lateral/axial PSFs and the co-registration across the FOV using fluorescent microspheres and report these metrics alongside any related image analysis, and—when object scale relative to the PSF (α) is known or estimated—apply the closed-form corrections or re- project to a declared reference PSF size/co-registration level. Where α is not directly available, brief auxiliary calibrations or transparent computational surrogates could suffice. In this manner, QC-aware normalization turns instrument variability from a confound into a controllable variable, improving interpretability, reproducibility, and comparability of fluorescence microscopy data at scale.

Finally, we note that the principles of this QC normalization strategy also align closely with the “reverse-thinking” framework proposed by Wait *et al*. for hypothesis-driven quantitative microscopy^79^. Their recommended approach for using light microscopy in research emphasizes beginning with a clearly articulated hypothesis, defining the specific data needed to test it, and then determining the experimental parameters and microscope configuration required to generate those data. By providing objective, quantitative metrics of instrument performance along with recorded image readouts, our suggested instrument normalization practice fits naturally within the experimental design phase of this framework and enables investigators to select imaging modalities and acquisition/analysis settings that are properly matched to the analytical image measurements required to test their hypotheses. Moreover, the central insight of this work is that image-derived biological measurements are not independent of instrument performance; rather, they are inherently constrained by measurable microscope QC parameters. Here, we have explicitly demonstrated these relationships for distance-based image morphometry and inter-channel overlap and colocalization measurements, but the concept is general. Many common quantitative image measurements can, in principle, be linked mathematically back to a corresponding or related opto-mechanical microscope performance QC parameter. Making these dependencies explicit provides a experimental mindset that builds performance validation into experimental design, so biological conclusions rely on measured, instead of assumed, imaging capability.

---

*Supplementary Fig. 1*

*Supplementary Fig. 2*

*Supplementary Fig. 3*

*Supplementary Fig. 4*

*Supplementary Fig. 5*

