## Supplementary Text for "Quality-control Normalization of Fluorescence Microscopy Morphometry and Colocalization Measurements for Improved Accuracy and Cross-instrument Reproducibility"

According to the wave theory of light, fluorescence emission is treated as an incoherent photophysical process. Individual excited fluorophores radiate spontaneously with mutually random phases, so interference cross terms average away and *intensities* (not complex fields) simply add together directly[1]. Consequently, within an isoplanatic region of the FOV, a fluorescence microscope behaves as a linear, shift-invariant system where the output image is the sum of identical point-spread functions (PSFs) shifted to each emitter's position—or equivalently, the object's fluorophore distribution convolved with the PSF—yielding a blurred representation of the underlying labeled structure[2]. For a diffraction-limited objective lens with a uniformly filled circular pupil, this intensity PSF is an Airy function (the squared magnitude of the Fraunhofer diffraction pattern of the pupil), which sets the fundamental resolution limit of the microscope[3]. In practice, the central lobe of the Airy function (containing most of the energy) is well approximated by a 2D Gaussian function (or by a Gaussian ellipsoid function in 3D) since the weaker secondary rings of the Airy function are often diminished or obscured by instrumental noise[4,5]. This Gaussian surrogate is analytically convenient: Gaussian convolutions remain Gaussian, so the net blur from multiple approximately Gaussian sources has a total variance equal to the sum of each source variance[6], letting us consider defocus, mild aberrational broadening, motion blur, and detector pixel digitization as simple additive contributions to resolution degradation. It also yields simple closed-form expressions that predict how blur and image mis-registration attenuate or amplify inter-channel pixel-wise correlation measurements like colocalization, volume overlap, etc. Thus, the incoherence of fluorescence lets us decompose the fluorescence microscope image into additive PSF building blocks, and the Gaussian approximation makes quantitative resolution and co-registration analyses algebraically tractable. Below we derive expressions for two common image measurements using these properties and show how the microscope's own QC metrics—the lateral/axial resolution and the channel co-registration—enter directly into and affect these quantities.

Volumetry of cellular biomolecules is one example of a common microscope image measurement used to probe and quantify various biological phenomena. In our context for this study, the volumes occupied by synaptic proteins in dendritic spines can, and often do serve as proxies to infer synapse connection strengths and maturation states under a variety of perturbations through so-called synaptogenesis image assays. Two-color versions of these image assays measure the volume overlap/co-occupation or intensity colocalization of protein signals within the AZ/PSD across the synaptic cleft to study neuronal growth and synapse connections as well. We first consider the volume of the microscope PSF itself. Since the PSF can be faithfully modelled as a Gaussian ellipsoid, the volumetric dimensions of this ellipsoid are

$$V_{\mathrm{PSF}} = \frac{4}{3}\pi\, abc \quad , \quad \text{(S1)}$$

With $a = b = \dfrac{\mathrm{FWHM}_{xy}}{2}$ and $c = \dfrac{\mathrm{FWHM}_{z}}{2}$

where $\mathrm{FWHM}_{xy}$ and $\mathrm{FWHM}_{z}$ refer to the measured lateral (assumed to be radially symmetric) and axial full-width at half-maximum extents of the Gaussian ellipsoid respectfully[7,8]. And so, the segmented volume of the PSF can be written as,

$$V_{\mathrm{PSF}} = \frac{\pi}{6}\mathrm{FWHM}_{xy}^{2} \cdot \mathrm{FWHM}_{z} \quad . \quad \text{(S2)}$$

Therefore, fluorescent objects (individual fluorescent proteins, molecular complexes, nanoscopic vesicles, etc.) in the specimen significantly smaller than the PSF lateral and axial extents will occupy a 3D image volume that matches this size because of diffraction. The volumes of fluorescent objects or sample features that are roughly equal to or physically larger than the PSF are blurred and can be calculated as follows: The 3D image intensity distribution of an incoherently emitting fluorescent object, $I(\mathbf{r})$, is a convolution of the object's 3D fluorescence distribution, $f_{obj}(\mathbf{r})$, with the microscope's 3D PSF intensity distribution, $p(\mathbf{r})$:

$$I(\mathbf{r}) = (f_{obj} * p)(\mathbf{r}) = \int_V f_{obj}(\mathbf{r}')\, p(\mathbf{r} - \mathbf{r}')\, d^3\mathbf{r}' \quad . \quad \text{(S3)}$$

if we consider a 1-dimensional Gaussian fluorescence distribution $f_{obj}(x)$ with variance term

$\sigma^2_{obj}$ ($\sigma = \sqrt{Var}$), this is:

$$f_{obj}(x) = \frac{1}{\sqrt{2\pi\sigma^2_{obj}}} \exp\left[-\frac{x^2}{2\sigma^2_{obj}}\right]. \qquad \text{(S4)}$$

The Fourier transform of a Gaussian function is also a Gaussian function[3], $\hat{f}_{obj}(k)$ with spatial frequency $k = \frac{1}{x}$:

$$\hat{f}_{obj}(k) = \exp\left[-(k\sigma_{obj})^2/2\right]. \qquad \text{(S5)}$$

And since convolution in real space is equivalent to multiplication in Fourier space (the convolution theorem), the Fourier transform of the image light intensity distribution, $\hat{I}(k)$, is:

$$\hat{I}(k) = \hat{f}_{obj}(k)\,\hat{p}(k) = \exp\left[-\tfrac{1}{2}k^2\left(\sigma^2_{\text{obj}} + \sigma^2_{\text{PSF}}\right)\right]. \qquad \text{(S6)}$$

The convolution of the sample object and the PSF (Equation S3) is computed through the inverse Fourier transform and shows that $I(x)$ is itself also Gaussian with a measured variance, $\sigma^2_{\text{meas}}$, equal to the sum of the fluorescence distribution and PSF distribution variances, which is known as variance addition (or standard deviation addition in quadrature):

$$I(x) = \frac{1}{\sqrt{2\pi(\sigma^2_{\text{obj}} + \sigma^2_{\text{PSF}})}} \exp\left[-\frac{x^2}{2(\sigma^2_{\text{obj}} + \sigma^2_{\text{PSF}})}\right] = \frac{1}{\sqrt{2\pi\sigma^2_{\text{meas}}}} \exp\left[-\frac{x^2}{2\sigma^2_{\text{meas}}}\right]. \qquad \text{(S7)}$$

For Gaussian functions, the FWHM is related to $\sigma$ through the identity:

$$\text{FWHM} = 2\sqrt{2ln2}\sigma \approx 2.355\sigma, \qquad \text{(S8)}$$

And therefore, along any Cartesian coordinate, the FWHMs add in quadrature:

$$\text{FWHM}^2_{meas} = \text{FWHM}^2_{obj} + \text{FWHM}^2_{PSF}. \qquad \text{(S9)}$$

With this relationship in hand, we can recognize then that small, isolated image features which are close in size to the PSF will also have a Gaussian ellipsoid-shaped volume, $V_{\text{meas}}$, that measures as:

$$V_{\text{meas}} = \frac{\pi}{6} \cdot [(\text{FWHM}_{\text{xy,obj}})^2 + (\text{FWHM}_{\text{xy,PSF}})^2] \sqrt{[(FWHM_{z,obj})^2 + (\text{FWHM}_{z,PSF})]^2} \quad . \text{(S10)}$$

From Equation S10, we can see that as the isolated object's fluorescence distribution extent uniformly grows in size, the PSF blurring contributions to the overall volume become negligible, and the measured volume approaches the true fluorescent object distribution volume. This behavior is observed in the simulated images of Supplementary Fig. 3 where the volumes of 1 um diameter and 2 um diameter fluorescent microspheres remain close to their true values even though the resolution degrades axially by a factor of 1.5 (and laterally by a factor of$\sqrt{1.5}$ concurrently) when imaged with a high NA oil immersion objective lens. Smaller simulated microspheres, on the other hand, grow and inflate accordingly. An important corollary of this fact is that a microscope system's resolution (a QC metric) significantly affects and factors into intra-channel distance-based morphometry measurements (length, area, volume, cluster size, etc.) *when the size of the measured sample feature is comparable to, or the same order of magnitude as, the resolution*. This is the reason why measurements of small objects from a microscope image only make sense in the context of the system resolution. As was shown in Fig. 8 and Fig. 9, the exact same presynaptic puncta were determined to have significantly different volumes depending on which objective lens was used to image them!

For the positive control resolution experiment, we measured reductions to the lateral and axial resolution resulting from closure of the microscope's emission pathway iris using a QC sample (microspheres) along with the accompanying volume inflations of small dendrite puncta in hippocampal neurons labeled by fluorescent active zone vGLUT proteins under the same iris conditions. Equation S10 allows us to write down a straightforward expression that directly links these QC calibrations to biological sample volumetric measurements taken under the different iris settings:

$$\frac{V_{\text{res}}}{V_{\text{full}}} = \frac{1 + \text{FWHM}^2_{xy,PSFres}/\text{FWHM}^2_{xy,obj}}{1 + \text{FWHM}^2_{xy,PSFfull}/\text{FWHM}^2_{xy,obj}} \sqrt{\frac{1 + \text{FWHM}^2_{z,PSFres}/\text{FWHM}^2_{z,obj}}{1 + \text{FWHM}^2_{z,PSFfull}/\text{FWHM}^2_{z,obj}}} \quad , \qquad \text{(S11)}$$

where the subscripts *res* and *full* refer to the restricted and full NA conditions respectively. This equation, combined with Equation S10 to determine the puncta extents (assuming a spherical fluorescence distribution such that these values were equal, ie: $\text{FWHM}_{xy,obj} = \text{FWHM}_{z,obj}$), was used to generate the solid black curve in Fig. 5b.

With this Gaussian PSF surrogate approximation, the same variance addition rule can be used to specify the measurement inflation factors associated with other common geometric quantities extracted from fluorescence microscope images. Since the measured FWHM along each axis is the quadrature sum of the true size and the PSF width, any intra-channel distance-based morphometric measurement inflates by the corresponding product of axis scale factors and can therefore be corrected for PSF blur. Let the true object FWHM along the three spatial axes be

$\mathrm{FWHM}_{\mathrm{obj},x}$, $\mathrm{FWHM}_{\mathrm{obj},y}$, $\mathrm{FWHM}_{\mathrm{obj},z}$ and the microscope PSF FWHMs be $\mathrm{FWHM}_{\mathrm{PSF},xy}$ (lateral) and $\mathrm{FWHM}_{\mathrm{PSF},z}$ (axial). Define the object-to-PSF ratios

$$\alpha_x = \mathrm{FWHM}_{obj,x}/\mathrm{FWHM}_{PSF,xy}, \quad \alpha_y = \mathrm{FWHM}_{obj,y}/\mathrm{FWHM}_{PSF,xy}, \quad \alpha_x = \mathrm{FWHM}_{obj,z}/\mathrm{FWHM}_{PSF,z}$$

and the axial-to-lateral PSF factor $k = \mathrm{FWHM}_{PSF,z}/\mathrm{FWHM}_{PSF,xy}$ (typically $k \approx$ 2-3 for high-NA systems). With FWHM quadrature addition, the measured (blurred) extents along each axis will be

$$\mathrm{FWHM}_{\mathrm{meas},x} = \mathrm{FWHM}_{PSF,xy}\sqrt{1+\alpha_x^2}, \quad \mathrm{FWHM}_{\mathrm{meas},y} = \mathrm{FWHM}_{PSF,xy}\sqrt{1+\alpha_y^2}, \quad \mathrm{FWHM}_{\mathrm{meas},z} = \mathrm{FWHM}_{PSF,z}\sqrt{k^2+\alpha_z^2}$$

These relations can be used in two equivalent ways: (i) Predicting the outcome of a measurement from known object size simply by plugging in the α values into the formulas above or, (ii) correcting measurements (“deflating” the blur) to estimate object size by inverting the equations:

$$\mathrm{FWHM}_{obj,x} = \sqrt{\mathrm{FWHM}_{\mathrm{meas},x}^2 - \mathrm{FWHM}_{PSF,xy}^2}$$

$$\mathrm{FWHM}_{obj,y} = \sqrt{\mathrm{FWHM}_{\mathrm{meas},y}^2 - \mathrm{FWHM}_{PSF,xy}^2}$$

$$\mathrm{FWHM}_{obj,z} = \sqrt{\mathrm{FWHM}_{\mathrm{meas},z}^2 - \mathrm{FWHM}_{PSF,z}^2}$$

For composite morphometric image measurements that are built from axis lengths (e.g., length, area, volume), the corrections propagate multiplicatively axis-by-axis. In PSF-normalized form, the “inflation factors” are:

- Length in $x$: $\sqrt{1+\alpha_x^2}$
- $xy$ Area: $\sqrt{1+\alpha_x^2}\,\sqrt{1+\alpha_y^2}$
- $xyz$ Volume: $\sqrt{1+\alpha_x^2}\,\sqrt{1+\alpha_y^2}\,\sqrt{k^2+\alpha_z^2}$

Therefore, to obtain a PSF-corrected estimate of a length/area/volume from measured data, replace each measured axis by its deflated form above and then insert it into the appropriate metric definition. The same process holds for other morphometry measurements like surface area, cluster size, etc. Plots of the inflation factors show that once α ≳ 2 the PSF contributes ≤25% error to measurements of volume, whereas diffraction-limited objects (α ≈ 1) can have volumes inflated by >2×.

Next, we consider how a microscope’s QC chromatic co-registration ratio factors into inter-channel multiplicative image measurements involving image intensity overlap (over areas, volumes, etc.) or covariance (colocalization). For simplicity, we start with a 1D version of the light intensity distribution of a small fluorescent object close in size to the diffraction limit

(Equation S8) in one channel—again leveraging the simpler mathematics of Gaussian approximations of fluorescence imaging—and consider its relationship to a second intensity distribution in another channel but displaced from the first channel object by some inter-channel distance, $r$, due to channel mis-registration. We will build to the more general 3D case thereafter.

Computation of the Hadamard intensity product between these two spectral objects (the basic calculation driving overlap and covariance-like image measurements), $\mathcal{I}_{meas}$, is accomplished through evaluation of the integral:

$$\mathcal{I}_{meas} = \int_{-\infty}^{\infty} I_1(x) I_2(x-r)\,dx = \mathcal{I}_0 \int_{-\infty}^{\infty} e^{-x^2/2\sigma^2_{1,meas}}\, e^{-(x-r)^2/2\sigma^2_{2,meas}}\,dx = \mathcal{I}_0\, e^{-r^2/2(\sigma^2_{1,meas}+\sigma^2_{2,meas})} . \quad \text{(S12)}$$

The subscripts *1* and *2* in Equation S12 refer to the channel number. The leading constant term, $\mathcal{I}_0$, of this equation is the “true” perfect image registration overlap intensity ($r = 0$), while the exponential term is a dampening factor that results from the channel mis-registration. As we will see below, this exponentially dampening (or amplification) of inter-channel overlap/covariance measurements is a general consequence of image misregistration, and this characteristic is retained in the 3D case. Therefore, even fractional displacements of one channel image with respect to another can lead to dramatically different measurement outcomes!

Faklaris *et al.*[7] define a microscope QC metric known as the system’s inter-channel or chromatic displacement index (CDI) co-registration ratio, which we denote here as $\chi$:

$$\chi = \frac{r}{r_{ref}} . \qquad \text{(S13)}$$

$\chi$ is the QC metric we extract from a two-channel, full FOV Z-scan of multi-color emitting fluorescent microspheres. $r_{ref}$ is the reference surface normalizing distance for the channel displacement, with the reference surface originally specified as the 3D ellipsoid envelope bounded by the *theoretical* resolution limits of the PSF associated with the shorter emission wavelength channel (see Equation S2 again)[9]. In 3D, $r_{ref}$ must be computed along the direction corresponding to the local channel displacement vector $\mathbf{r}$. In the resolution-volume morphometry experiments of this study, we observed three distinctly different lateral and axial resolutions associated with the PSF of each objective lens. As stated in the main text, we suggest a more realistic calculation of $r_{ref}$ that should be based on *real, measured* FWHM resolutions instead of theoretical limits, as this will lead to more accurate determination of co-registration ratios. Supplementary Fig. 5 displays the differences in the resulting $\chi$ heatmaps and axial vs. lateral channel displacement scatterplots when these different normalizations are used.

In our 1D treatment of the intensity overlap between two small Gaussian-like fluorophore distributions, $r_{ref}$ is simply the measured 1D PSF lateral extent for the shorter wavelength Channel 1, $FWHM_{x,PSF,1}$:

$$\chi = \frac{r}{FWHM_{x,PSF,1}}. \quad \text{(S14)}$$

Thus, when this normalization of the inter-channel mis-registration distance is combined with Equations S8 and S9, Equation S12 can be re-written in terms of $\chi$ and measured channel FWHMs as:

$$\mathcal{I}_{meas} = \mathcal{I}_0 \exp\left[-4ln2\chi^2 \frac{\mathrm{FWHM}^2_{\mathrm{x,PSF,1}}}{\mathrm{FWHM}^2_{x,meas,1} + \mathrm{FWHM}^2_{x,meas,2}}\right]. \quad \text{(S15)}$$

Note that when there is no channel mis-registration ($\chi$=0), the true inter-channel intensity overlap signal between the two objects, $\mathcal{I}_0$, is again recovered. If the imaged object fluorescent intensity distributions are the same in both channels ($\mathrm{FWHM}_{x,meas,1} = \mathrm{FWHM}_{x,meas,2}$), and if the PSF size differences due to emission wavelength or channel aberrations are negligible ($\mathrm{FWHM}_{x,PSF,1} = \mathrm{FWHM}_{x,PSF,2}$), then Equation S15 takes a simplified form of:

$$\mathcal{I}_{meas} = \mathcal{I}_0 \exp\left[-\frac{2ln2}{1+\alpha^2}\chi^2\right], \quad \text{(S16)}$$

with $\mathrm{FWHM}^2_{x,meas} = \mathrm{FWHM}^2_{x,PSF}(1+\alpha^2)$,

and $\alpha = \frac{FWHM_{x,obj}}{FWHM_{x,PSF}}$,

which is just Equation S9 rearranged into a convenient size factor ratio. There are a couple noteworthy characteristics of expression S16:

1. When the object extent is much smaller than the extent of the PSF ($\alpha \ll 1$), the exponential term reduces to $e^{-2ln2\chi^2}$. As we shall see below, this exponential factor is general and also appears in the 3D treatment as well. Consider the value of $\chi$=1, which is the upper acceptable tolerance for channel mis-registration recommended by Faklaris *et al.* and the QUAREP-LiMi consortium[7,10]. This corresponds to the scenario where the center of Channel 2's PSF lies exactly on the reference surface FWHM envelope of Channel 1's PSF. At this tolerance level, any inter-channel overlap or colocalization image measurement of a diffraction-limited sized co-labeled object will be reduced by 75% and subject to the noise levels on the measurement signal. Thus, the

recommended tolerance level of $\chi$=1 is somewhat arbitrary without justification. The measurement precision can be improved simply by confining the image measurement to smaller regions or areas of the FOV where the channel mis-registration is known to be less by means of a QC test.

2. When the object's extent is very large compared to the PSF extent (large $\alpha$), the exponential term reduces to 1, and just as we saw with our resolution study (and intuitively), there is a certain size of an imaged object, relative to channel mis-registration separation distance, where the mis-registration has little impact on any inter-channel measurement of overlap or covariance.

Continuing with our 1D treatment of the situation, let us now consider the case where the fluorescent objects in channels 1 and 2 do not occupy the same physical location and instead are separated by some biologically relevant distance $\delta$ from one another, and let $\tilde{\delta} = \delta / FWHM_{x,PSF}$. Under this condition, the integral of Equation S12 becomes:

$$\mathcal{I}_{meas} = \mathcal{I}_0 \exp\left[-\frac{2ln2}{1+\alpha^2}\left(\chi - \tilde{\delta}\right)^2\right]. \qquad \text{(S17)}$$

The additional mixed term proportional to $-2\tilde{\delta}\chi$ in the exponential can either further dampen or inflate the inter-channel overlap measurement depending on the sign and magnitude of $\delta$ (object separation can be along the channel mis-registration direction $r$ or opposite to it).

Extension of this model into three dimensions hinges on the fact that if the PSFs of both channels are axis-aligned with each other, then the 3D version of Equation S12 factorizes into three separable 1D Gaussian integrals. It is reasonable to assume co-aligned channel PSFs since the excitation and detection optical pathways are rotationally symmetric around the z-axis (the microscope optic axis) and the camera defines the orthogonal x- and y-axes. The dominant PSF anisotropy is axial versus lateral widths, with the principal PSF ellipsoid axes aligned to (x, y, z). Tilted PSFs can arise from aberrations that couple system axes together (e.g., astigmatism at an oblique angle, coma, etc.) or from far-off z-axis field positions. In this study, we did not observe aberrant or tilted inter-channel PSF alignment of this nature, and so we ignore the possibility of this complication to the analysis.

We define the net separation vector (biology + channel mis-registration), $\boldsymbol{\Delta}$, as:

$$\boldsymbol{\Delta} = \boldsymbol{\delta} + \mathbf{r}, \qquad \text{(S18)}$$

or in Cartesian form $\boldsymbol{\Delta} = (\Delta_x, \Delta_y, \Delta_z)$. First, we normalize the components of $\boldsymbol{\Delta}$ by the shorter wavelength channel PSF FWHMs in a similar fashion as for the 1D case to obtain the vectorized components of the cross-channel co-registration:

$$\chi_x = \frac{\Delta_x}{FWHM_{x,PSF}}, \quad \chi_y = \frac{\Delta_y}{FWHM_{y,PSF}}, \quad \chi_z = \frac{\Delta_z}{FWHM_{z,PSF}}. \tag{S19}$$

And define per-axis object to PSF size factors:

$$\alpha_x = \frac{FWHM_{x,obj}}{FWHM_{x,PSF}}, \quad \alpha_y = \frac{FWHM_{y,obj}}{FWHM_{y,PSF}}, \quad \alpha_z = \frac{FWHM_{z,obj}}{FWHM_{z,PSF}}. \tag{S20}$$

The 3D form of Equation S17 is then:

$$\mathcal{I}_{meas\,3D} = I_0 \exp\left\{-2\ln 2\left[\frac{(\chi_x - \bar{\delta}_x)^2}{1+\alpha_x^2} + \frac{(\chi_y - \bar{\delta}_y)^2}{1+\alpha_y^2} + \frac{(\chi_z - \bar{\delta}_z)^2}{1+\alpha_z^2}\right]\right\}. \tag{S21}$$

Just as we saw a direct relationship between intra-channel volumetric measurements and channel resolution (Equation S10), here too in Equation S21 we observe a direct relationship between inter-channel overlap image measurements and a microscope QC metric (the channel co-registration, $\chi$), as well as the channel resolution and its relative size to the imaged object again via the $\alpha_i$ terms. The 3D form of this relationship emphasizes that prediction (or correction) of intensity overlap or colocalization attenuation/amplification depends on knowledge of the channel mis-registration orientation with respect to the biologically relevant separation distance vectors and potentially anisotropic structural orientations of the objects in both channels. The channel mis-registration dampening/amplification effect is large when channel shifts align with thin fluorescent object directions (small $\alpha_i$) and small when along extended directions (large $\alpha_i$). Co-labeled fluorescent microtubules are a good example of both scenarios. Thin sheets or membranes are broad in xy, but thin in z. A small axial mis-registration can dampen the image measurement appreciably, while lateral mis-registration may matter less. In the general case, sensitivity to mis-registration is controlled by how much fine detail (small correlation length) the structure has along the direction of the shift after PSF blur. When a structure is large and smooth relative to both the PSF and the shift, the damping becomes negligible; when it is thin or finely textured along any axis, the damping remains measurable, and it is strongest for shift components across that thin dimension. The anisotropic formula above captures this conceptualization in one expression.

Another way to view the general case is through the Fourier domain: Channel mis-registration mainly suppresses the high spatial frequency content of the inter-channel product. If a sample's structure has little high-frequency content along a certain direction, registration errors along that direction impart little effect onto the image measurement. In contrast, if the sample structure does have high-frequency content along a certain direction (thin or textured), registration errors along that direction play a bigger role in the resulting outcome of the measurement.

A single scalar co-registration collapses direction into one number. Two ROIs can have the same scalar $\chi$ but very different directional decompositions, leading to different damping. The orientation of channel mis-registration **r** relative to the imaged object's fine structure is key, and the scalar χ hides that geometry. In other words, a microscope's co-registration QC metric would best be reported as a full 3D *vector* map (directionally-resolved) across the entire FOV as opposed to a *scalar* map like we did in Fig. 10a for each objective lens. Nevertheless, the scalar maps of channel co-registration are convenient and easier to work with, and they still hold utility in terms of microscope QC and experimental design.

For isolated, punctate sample features where the biological structure is randomly oriented in space (isotropic) and when the imaged region contains an ensemble of labeled structures in both channels, the total $\boldsymbol{\delta}$ vector averages to zero ($\langle \delta_{tot} \rangle \approx 0$) and the $\boldsymbol{\Delta}$ vector becomes just $\mathbf{r}$. The same thing occurs when there is true biological colocalization ($\boldsymbol{\delta}$=0). In our study, we assume any given pre/post synaptic vector imaged across both channels is randomly oriented. A special case of a two-channel imaging assay permitting application of Equation S21 in a reduced scalar form occurs when we further assume that the imaged objects have roughly equal object-to-PSF size ratios in all 3 dimensions ($\alpha_x \approx \alpha_y \approx \alpha_z = \alpha$) and in both channels. Recognizing that:

$$\chi^2 = \chi_x^2 + \chi_y^2 + \chi_z^2, \qquad \text{(S22)}$$

allows us to rewrite Equation S21 as:

$$\mathcal{I}_{meas\,3D} = \mathcal{I}_0 \exp\left[-\frac{2ln2}{1+\alpha^2}\,\chi^2\right], \quad \text{(S23)}$$

which matches Equation S16.

In Fig. 10a, we mapped the local *scalar* Green-to-Yellow inter-channel co-registration $\chi$ value across the FOV for Objectives 1-3 and showed that each objective possessed a zone of maximal co-registration (not always at the FOV center - see Objective 2) which decayed radially outwards from this zone. With this map, it is possible to predict how much $\mathcal{I}_{meas\,3D}$ will vary as a dual-labeled sample image feature is translated from one region of interest (ROI1) in the FOV to another (ROI2), provided that the PSF is spatially invariant across the FOV in both channels. This latter point is validated in our study by the observation that the PSFj resolution heatmaps were relatively flat for all three objective lenses (Fig. 7a). Under these conditions, the inter-channel overlap/covariance signal change ratio between the two FOV regions of interest, $\mathcal{I}_{meas\,3D,ROI2}/\mathcal{I}_{meas\,3D,ROI1}$, is:

$$\frac{\mathcal{I}_{meas\,3D,ROI2}}{\mathcal{I}_{meas\,3D,ROI1}} = \exp\left[-\frac{2\ln 2}{1+\alpha^2}\left(\chi_{ROI2}^2 - \chi_{ROI1}^2\right)\right]. \quad \text{(S24)}$$

In the main text of this publication, we use Equation S24 to show how the measured colocalization signal between fluorescent vGLUT and Homer pre- and postsynaptic puncta predictably decays according to the spatial differences in co-registration which were inferred from the microsphere QC experiments for each objective lens. Equation S24 also demonstrates the somewhat counterintuitive tradeoff between microscope resolution and chromatic co-registration spatial dependencies: A microscope with degraded resolution (larger $\alpha$) is less sensitive to co-registration differences, whereas sharper resolutions increase sensitivity to these spatial changes in channel registration. Knowledge of both of these critical instrument parameters can therefore be used to fine-tune experimental precision and tailor the image acquisition location and area to a size that is optimal or sufficient for the biological process under examination. Alternatively, Equation S24 can be applied to error correct recordings of puncta colocalization across the FOV.

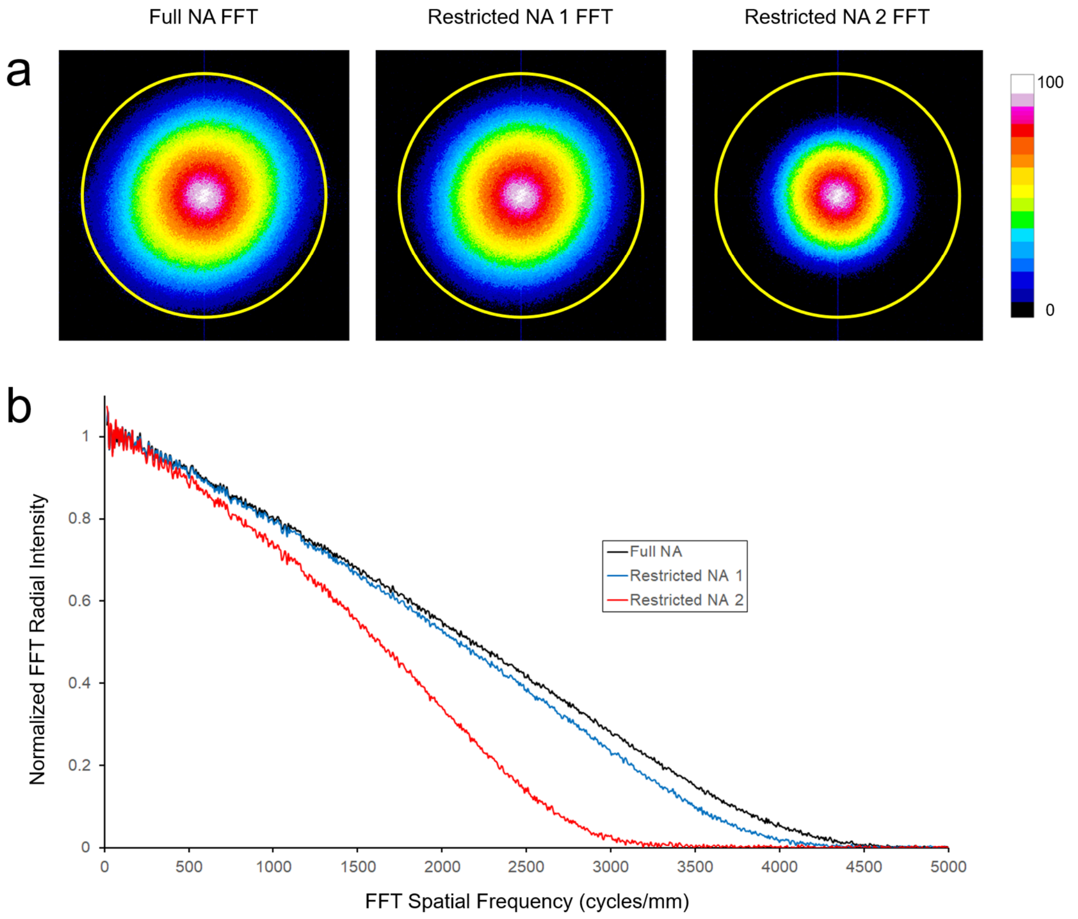


c

| | Full NA | Restricted NA 1 | Resticted NA 2 |
|---|---|---|---|
| MTF cut-off frequency (cycles/mm) | 4447 | 4193 | 3197 |
| MTF-derived Lateral Resolution (nm) | 225 | 239 | 313 |
| MTF-derived Lateral Resolution (Normalized) | 1.18 | 1.26 | 1.65 |
| MTF-derived Lateral Resolution %Change | | 6 | 28 |

**Supplementary Fig. 1 | Full FOV microscope lateral resolution characterization under varying detection NA conditions using Fourier domain analysis techniques. a**, The fast Fourier Transform (FFT) of the maximum intensity Z-projection images of microspheres under different detection NA settings of the control experiment were created in the open-source image analysis program ImageJ[11] and normalized to their peak intensities, producing nearly circularly symmetric spatial frequency spectrum distribution maps (also known as the 2D amplitude optical transfer function or modulation transfer function - MTF). **b**, The spatial frequency distribution maps were radially averaged using the Radial Profile Plot ImageJ plugin[12] and show progressively decreasing MTF bandwidths as the emission pathway iris was closed down. **c**, The 1% intensity level in the tail regions of each radially average spectrum curve was used to determine the MTF cut-off frequency in cycles/mm and back-calculated into spatial domain lateral resolutions. The MTF-derived resolutions and resolution percent differences compared to the Full NA condition are similar in magnitude to, but deviate from those determined by spatial domain PSF analysis techniques via PSFj (Fig. 3b). These differences could be a result of the difficulty in accurately locating the 1% cut-off spatial frequency and the arbitrariness in selection of the cut-off frequency level (1%, 5%, etc.). We chose 1% as a cut-off level in this case because of the high fidelity of the MTF spectrum curves. This technique, which interrogates the low signal tail regions of each spectrum is also extremely sensitive to the noise due to finite pixel size sampling and the size of the test object (200 nm microspheres) that are not quite below the diffraction limit for this objective lens (ie: not a true delta function). Together, these additional factors likely account for the discrepancies in resolution changes assessed between the spatial and frequency domain analysis approaches.

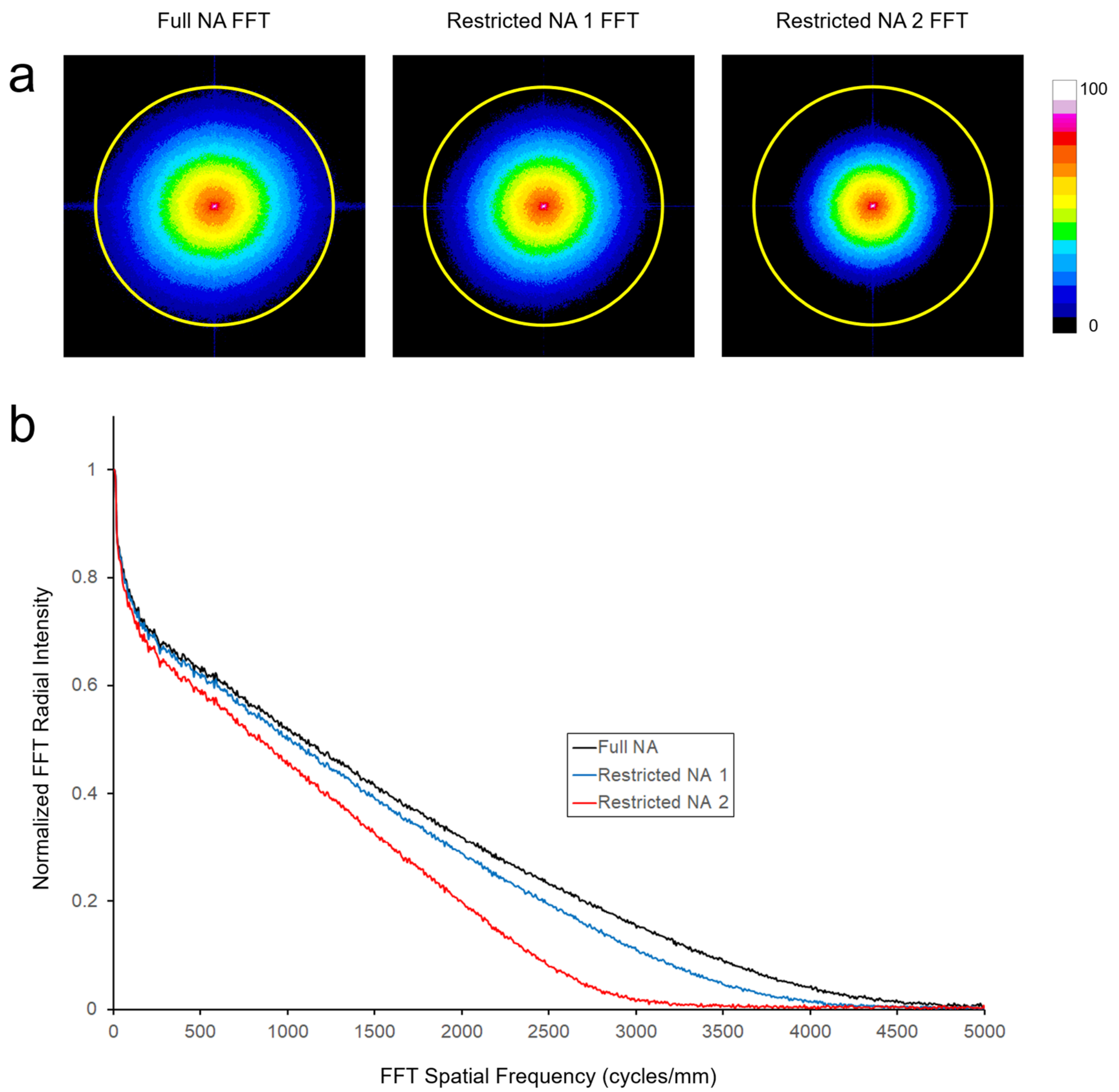


**Supplementary Fig. 2** | **Fourier-domain analysis of cultured neuron images under varying emission pathway NA conditions (matched to the microsphere QC experiment). a**, Maximum-intensity Z-projections from the same FOV of neurons expressing fluorescently tagged vGLUT proteins were Fourier transformed. The resulting 2D spatial frequency maps were normalized to unity at zero spatial frequency for each NA condition. **b**, Radial averages of these maps yield 1D MTF curves. Stopping down the emission pathway iris (Restricted NA 1, Restricted NA 2) narrows the spectra and lowers the lateral cutoff frequency relative to Full NA, indicating increased blur and reduced effective resolution, consistent with Supplementary Fig. 1.

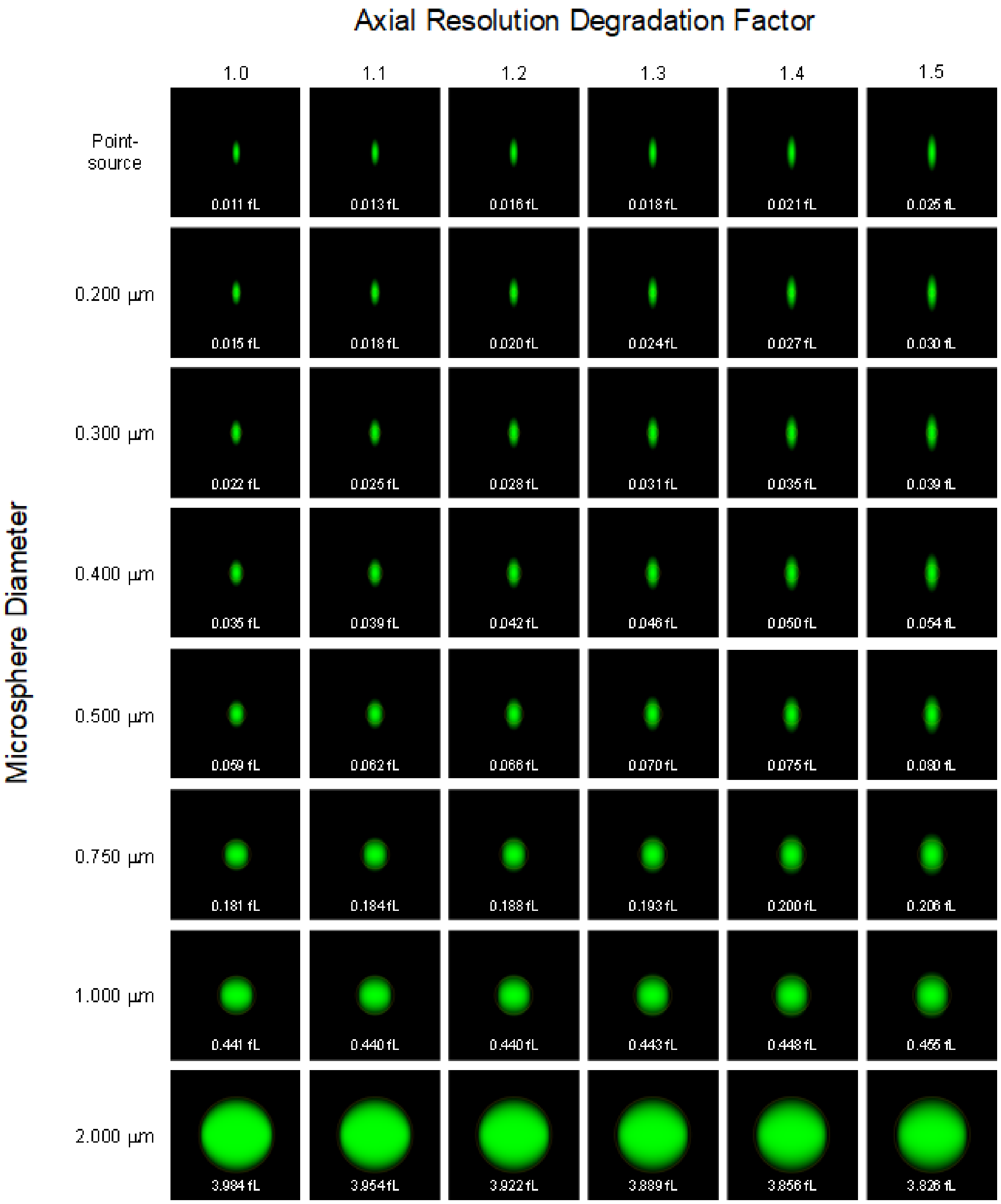


**Supplementary Fig. 3 |Simulations of different sized fluorescent microspheres and their apparent volumes when viewed under varying resolution conditions.** Each image row shows fluorescent spheres of gradually increasing sizes from an infinitely small (point source) up to a 2 µm sphere. Each column of images shows the apparent sphere's side-profile shape with decreasing effective NA of the system. The ideal system (Axial Resolution factor 1) represents a system with NA = 1.4. Each image lists the sphere's calculated volume (in fL). Once the spherical object becomes greater than 0.5 µm in diameter, there is very little difference in the apparent volume under changing resolution conditions (up to an axial resolution degradation factor of 1.5). What is perhaps less intuitive is that the object must be significantly larger than the theoretical resolution limit of the system before reaching this threshold.

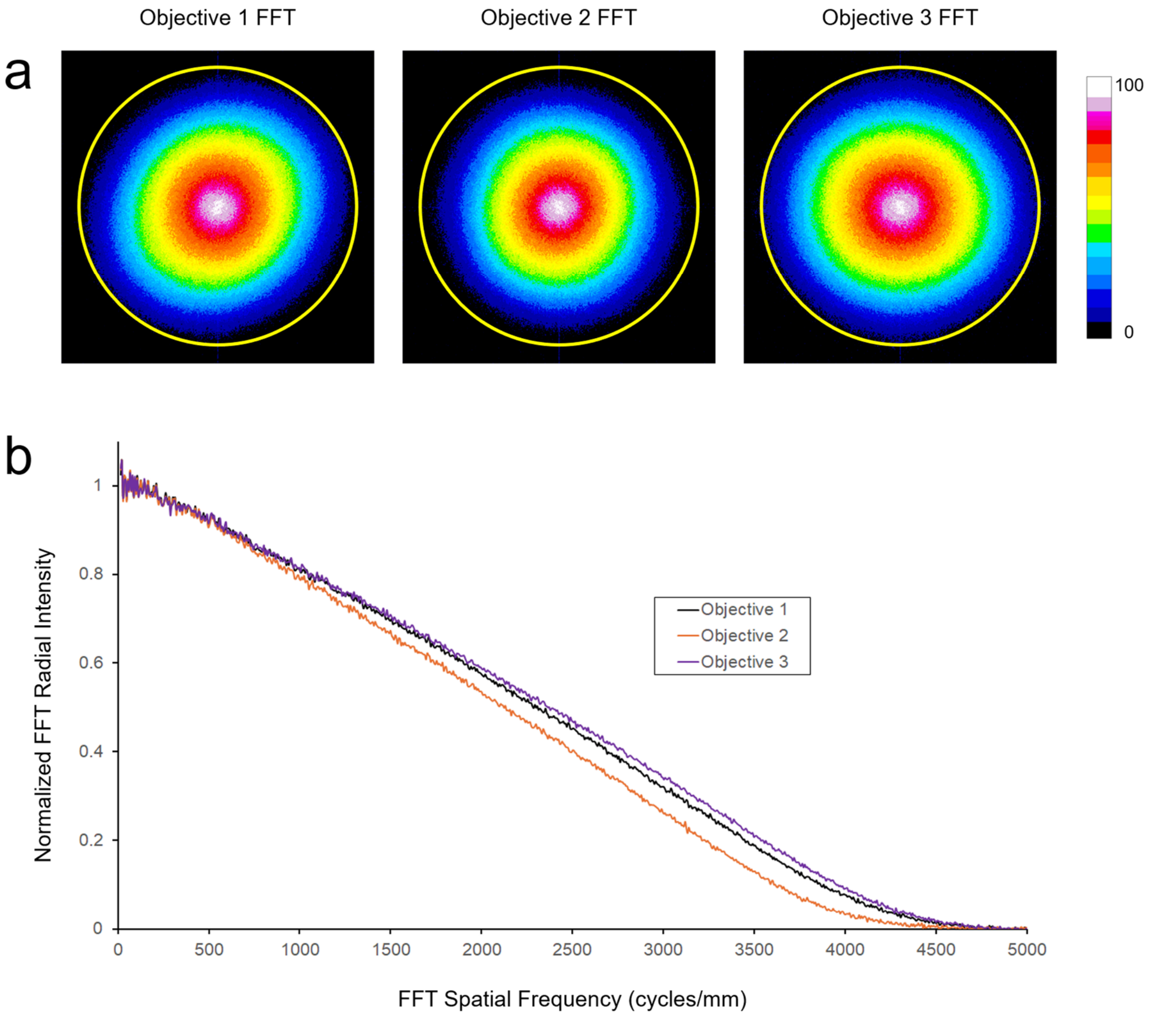


c

| | Objective 1 | Objective 2 | Objective 3 |
|---|---|---|---|
| MTF cut-off frequency (cycles/mm) | 4577 | 4310 | 4622 |
| MTF-derived Lateral Resolution (nm) | 218 | 232 | 216 |
| MTF-derived Lateral Resolution (Normalized) | 1.13 | 1.20 | 1.14 |

**Supplementary Fig. 4 | Full FOV microscope lateral resolution characterization of the Objective lenses 1-3 using Fourier domain analysis techniques**. **a**, The fast Fourier Transform (FFT) of the maximum intensity Z-projection images of the microspheres for Objectives 1, 2, and 3. **b**, Radially averaged MTFs for these objective lenses. **c**, MTF 1% level cut-off spatial frequencies and MTF-derived lateral resolutions (absolute and normalized to their theoretical limits) in tabular format.

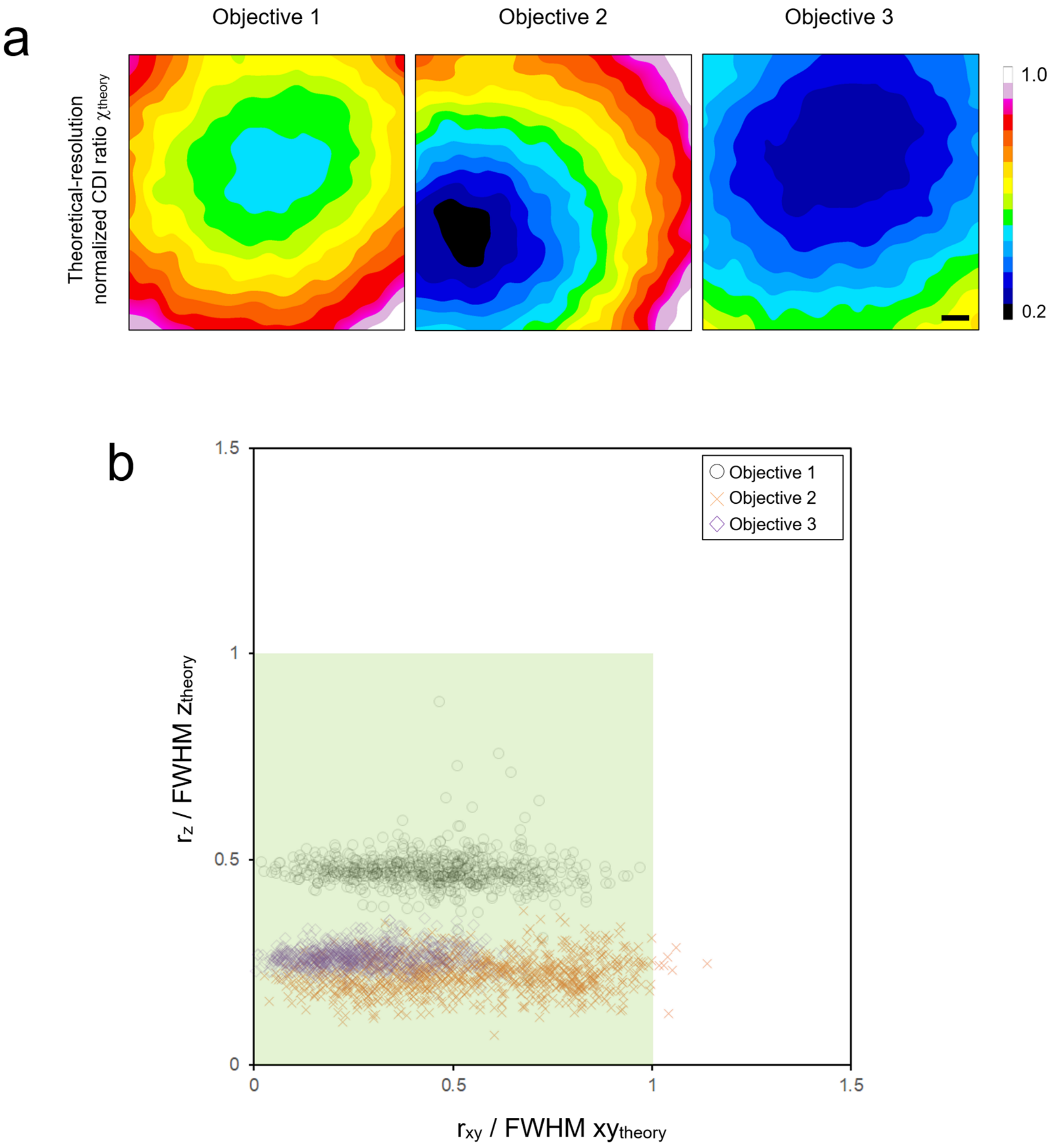


**Supplementary Fig. 5 | Full field of view objective lens chromatic co-registration characterization using theoretical PSF normalization. a**, Full-FOV CDI co-registration heatmaps computed by normalizing the measured channel displacements with the theoretical lateral and axial diffraction-limited FWHMs for the shorter-wavelength channel. One heatmap is shown per objective (Objectives 1–3). Scale bar = 20 µm. **b**, Scatterplot of axial vs. lateral normalized displacements for all identified microspheres. The green region marks the Faklaris acceptance window (χ < 1 in both axes).
